# ELOF1 is a transcription-coupled DNA repair factor that directs RNA polymerase II ubiquitylation

**DOI:** 10.1101/2021.02.25.432427

**Authors:** Yana van der Weegen, Klaas de Lint, Diana van den Heuvel, Yuka Nakazawa, Ishwarya V. Narayanan, Noud H.M. Klaassen, Annelotte P. Wondergem, Khashayar Roohollahi, Janne J.M. van Schie, Yuichiro Hara, Mats Ljungman, Tomoo Ogi, Rob M.F. Wolthuis, Martijn S. Luijsterburg

**Author notes:** These authors contributed equally to this work (Yana van der Weegen, Klaas de Lint).

## Abstract

Cells employ transcription-coupled repair (TCR) to eliminate transcription-blocking DNA lesions. The binding of the TCR-specific repair factor CSB triggers DNA damage-induced ubiquitylation of RNA polymerase II (RNAPII) at a single lysine (K1268) by the CRL4^CSA^ ubiquitin ligase. However, how the CRL4^CSA^ ligase is specifically directed toward the K1268 site is unknown. Here, we identify ELOF1 as the missing link that facilitates RNAPII ubiquitylation, a key signal for the assembly of downstream repair factors. This function requires its constitutive interaction with RNAPII close to the K1268 site, revealing ELOF1 as a specificity factor that positions CRL4^CSA^ for optimal RNAPII ubiquitylation. Furthermore, drug-genetic interaction screening reveals an unanticipated compensatory TCR pathway in which ELOF1 together with known factors DOT1L and HIRA protect CSB-deficient cells from collisions between transcription and replication machineries. Our study provides a genetic framework of the transcription stress response and reveals key insights into the molecular mechanism of TCR.

## Introduction

The transcription of protein-coding and non-coding genes involves RNA polymerase II enzymes (RNAPII), which synthesize RNA transcripts complementary to the DNA template strand. The presence of DNA lesions in the template strand causes stalling of elongating RNAPII leading to a genome-wide transcriptional arrest (Brueckner et al., 2007; Nakazawa et al., 2020; Tufegdžić Vidaković et al., 2020). It is essential that cells overcome this arrest and restore transcription to maintain gene expression and viability. The transcription-coupled nucleotide excision repair (TCR) pathway efficiently removes transcription-blocking DNA lesions through the CSB, CSA, and UVSSA proteins (Nakazawa et al., 2012; Schwertman et al., 2012; van der Weegen et al., 2020). Inactivating mutations in the *CSB* and *CSA* genes cause Cockayne syndrome, which is characterized by severe developmental and neurological dysfunction due to the prolonged arrest of RNAPII at DNA lesions (Laugel et al., 2010; Nakazawa et al., 2020).

The sequential and cooperative actions of CSB, CSA, and UVSSA recruit TFIIH to DNA damage-stalled RNA polymerase II to initiate DNA repair (van der Weegen et al., 2020). In addition to protein-protein contacts, TCR complex assembly is controlled by ubiquitylation. Our recent findings show that efficient transfer of TFIIH onto RNAPII requires ubiquitylation of a single lysine on the largest subunit of RNAPII (RPB1-K1268), which is essential for efficient TCR (Nakazawa et al., 2020). This DNA damage-induced modification of RNAPII is fully dependent on cullin-ring type E3-ligases (CRLs), and was found to be strongly decreased in CSA-deficient cells (Nakazawa et al., 2020), indicating that the CRL4^CSA^ E3 ligase complex is a critical regulator of RNAPII ubiquitylation.

The full repertoire of human factors required for TCR is currently unknown. The CSB protein, which may bind to DNA that emerges behind RNAPII right after transcription (Xu et al., 2017) (Fig. S1a), recruits the CRL4^CSA^ complex through an evolutionary conserved motif in its C-terminus (van der Weegen et al., 2020). The ligase arm in the CRL4^CSA^ complex is highly flexible and creates a defined ubiquitylation zone with CSA as its anchor point (Fischer et al., 2011). However, it remains to be elucidated how the CRL4^CSA^ ubiquitin ligase activity is specifically directed towards the K1268 site, which is located in front of RNAPII close to where the DNA enters the enzyme (Fig. S1b).

## Results

### A genome-wide CRISPR screen identifies *ELOF1* as a putative TCR gene

To identify unknown genes required for TCR, we performed a genome-wide CRISPR screen in the presence or absence of the natural compound Illudin S, which induces transcription-blocking DNA lesions that are selectively eliminated by TCR (Jaspers et al., 2002). For this CRISPR screen, we used the genome-wide pLCKO-TKOv3 library consisting of 71.090 exon-targeting sgRNAs (two component version of the library published in (Mair et al., 2019)). Three replicate pools of diploid, doxycycline-inducible Cas9-expressing RPE1-hTERT cells (hereafter called RPE1- iCas9), which were knockout for both *TP53* and the *PAC* puromycin-resistance gene, were transduced with the pLCKO-TKOv3 library. Transduced cells were selected with puromycin, treated with doxycycline to induce Cas9, and subsequently cultured for 12 population doublings in the presence or absence of up to 25 nM Illudin S. Genomic DNA was isolated, lentiviral inserts were amplified by PCR, sgRNA sequences were identified by next-generation sequencing (NGS), and the results were analyzed for enrichment or depletion of sgRNAs using drug-Z (Colic et al., 2019) (Fig. 1a).

**Fig. 1.**
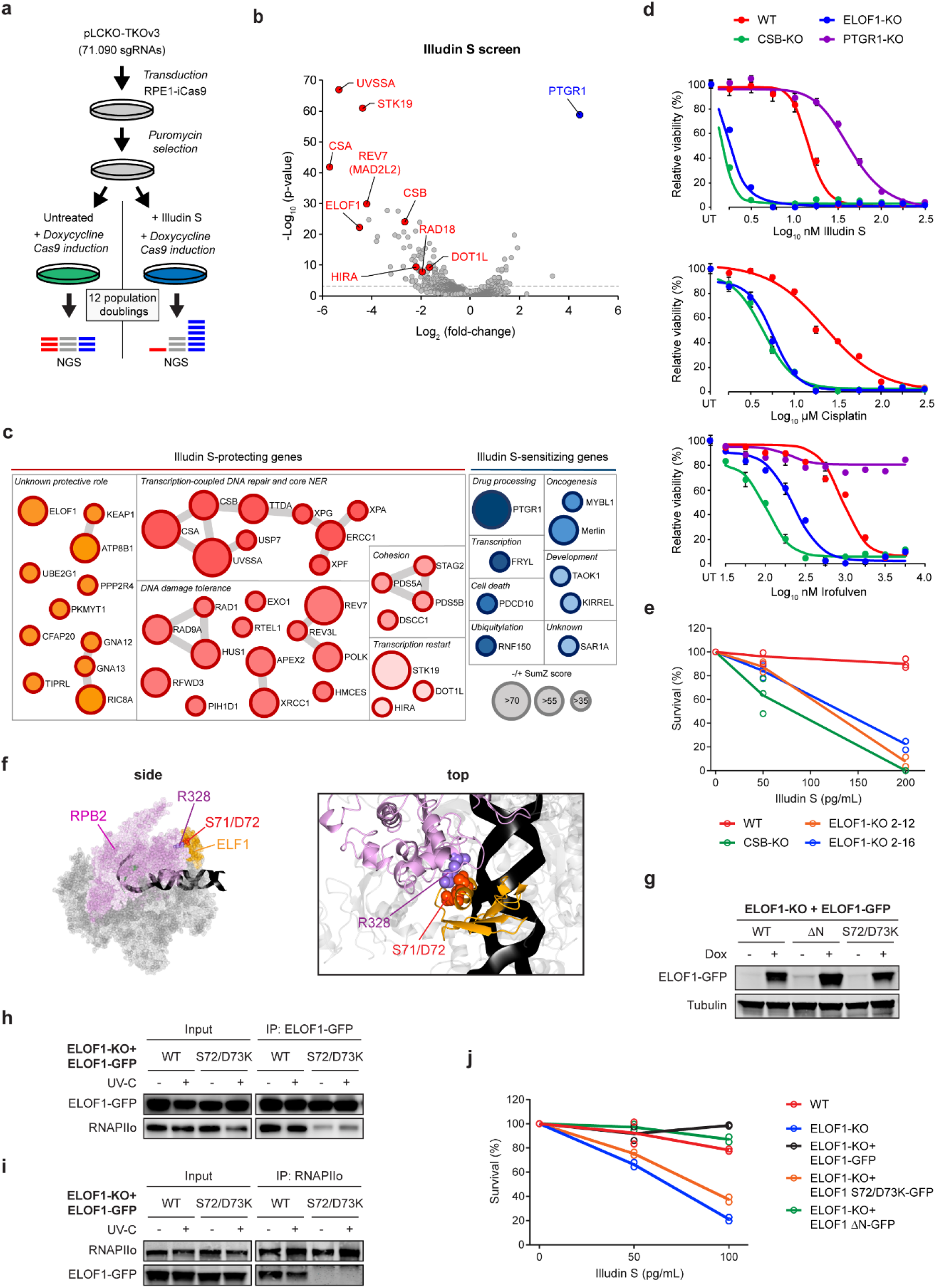
RNAPII-associated ELOF1 is a putative TCR gene. (**a**) Schematic representation of the CRISPR screening pipeline in RPE1-iCas9 cells set-up to identify new TCR genes. (**b**) Volcano plot depicting gene-knockouts sensitizing (selected genes highlighted in red) or conferring resistance (selected genes highlighted in blue) to Illudin S treatment (x-axis, log_2_ fold-change), and their significance (y-axis, -log_10_ p-value; full analysis results in Supplementary data). See Fig S2a for single guide counts. (**c**) Network analysis of highly significant hits representing genes essential for Illudin S resistance (red and orange) or promote Illudin S toxicity (blue). Grey lines reflect known protein-protein interactions (Cytoscape, BioGRID). (**d**) 72h drug sensitivity assays of indicated RPE1-iCas9 KO clones (*n=3*). Values indicate the average ± SEM of 6 technical replicates from one experiment (**e**) Clonogenic Illudin S survival of the indicated single RPE1- iCas9 KO clones. Each symbol represents the mean of an independent experiment (*n=3* for all except for *ELOF1*-KO 2-12 which is *n=2*), each containing 2 technical replicates. (**f**) Side-view and top-view (zoom) of the structure (PDB: 5XON) of *K. pastoralis* ELF1 (orange) bound to RNAPII (grey) with RPB2 in purple. Residues important for the ELF1-RNAPII interaction are indicated. (**g**) Western blot analysis of U2OS (FRT) *ELOF1*-KO cells complemented with inducible GFP-tagged versions of ELOF1. (**h**) Co-immunoprecipitation (IP) of ELOF1^WT^-GFP and ELOF1^S72/D73K^-GFP on the combined soluble and chromatin fraction (*n=4*). (**i**) Endogenous RNAPIIo Co-IP on U2OS (FRT) *ELOF1*-KO cells complemented with ELOF1^WT^-GFP and ELOF1^S72/D73K^-GFP (*n=4*). (**j**) Clonogenic Illudin S survival of U2OS (FRT) WT, *ELOF1*-KO, and GFP-tagged ELOF1 rescue cell lines. Each symbol represents the mean of an independent experiment (*n=2*), each containing 2 technical replicates.

Using a strict FDR cutoff of 0.01, we found 104 hits sensitizer hits and 18 hits conferring resistance to Illudin S (see supplemental analysis data). Strong resistance was conferred by gRNAs targeting *PTGR1* (Fig. 1b,c, Fig S2a), in line with the known role of the PTGR1 enzyme in bio- activating Illudin derivatives (Yu et al., 2012). Nine known core TCR genes, including *CSB, CSA* and *UVSSA* were identified, as well as several genes that were previously connected to transcription recovery after UV, including *HIRA* (Adam et al., 2013), *DOT1L* (Oksenych et al., 2013), and *STK19* (Boeing et al., 2016) (Fig. 1b,c, Fig S2a). Additionally, the 9-1-1 complex (Rad9-Hus1-Rad1) and several translesion synthesis components, including REV7 (MAD2L2) and RAD18 are required for cells to tolerate Illudin S, in line with the known ability of Illudin S-induced lesions to block replication (Jaspers et al., 2002) (Fig. 1b). These results validate the specificity and resolution of our CRISPR screening approach. In addition, our screen identified the previously uncharacterized *ELOF1* gene as a top hit, which we decided to study further (Fig. 1b, Fig S2a).

We generated single *PTGR1, ELOF1* and *CSB* knockouts in RPE1-iCas9 cells (Fig. S2b) and exposed the cells to compounds that generate transcription-blocking DNA lesions, such as Illudin S, its clinically used analogue Irofulven, and cisplatin. Drug-sensitivity assays confirmed that knocking out *ELOF1* causes strong sensitivity to transcription-blocking DNA damage, similar to TCR-deficient *CSB* knockout cells, while *PTGR1* knockouts were highly resistant to Illudin S and particularly Irofulven (Fig. 1d, Fig S3a). In addition, clonogenic Illudin S survival experiments showed that *ELOF1*-KO cells are nearly as sensitive to Illudin S as *CSB*-KO cells (Fig. 1e). Importantly, inducing the expression of a GFP-tagged ELOF1 in *ELOF1* knockout U2OS cells completely restored their Illudin S resistance (Fig. S3b-c). These findings identify ELOF1 as a promising candidate TCR gene.

### ELOF1 protects against transcription-blocking DNA damage by binding to RNAPII

ELOF1 is a small zinc-finger protein that is evolutionary conserved across eukaryotes, except for the C-terminal acidic tail (amino acids 84-145 in *S. cerevisiae*) which is absent in metazoan orthologues (Fig. S3d) (Daniels et al., 2009). ELF1, the yeast orthologue of ELOF1, is a putative transcription elongation factor that associates with RNAPII and promotes passage through nucleosomes together with the elongation factors SPT4, SPT5, and TFIIS (Ehara et al., 2019; Ehara et al., 2017). Yeast ELF1 binds between the RPB1 clamp-head domain and RPB2 lobe domain of RNAPII thereby preventing nucleosomes from being trapped between RPB1 and RPB2. Yeast ELF1 contains a central RNAPII-binding helix that interacts with RPB2 and an N-terminal basic tail that interacts with the incoming DNA (Fig. 1f, Fig S1c). Mutations in the RNAPII-binding helix (ELF1^S71/D72K^) or removal of the N-terminal DNA-binding region compromises the function of yeast ELF1 (Ehara et al., 2019; Ehara et al., 2017).

To directly assess whether these functions of ELOF1 are required to protect cells against transcription-blocking DNA damage, we stably expressed inducible versions of GFP-tagged ELOF1^WT^, ELOF1^Δ N^, and ELOF1^S72/D73K^ in *ELOF1*-KO cells (Fig. 1g). Pull-down of GFP-tagged ELOF1^WT^ showed a strong interaction with Ser2-phosphorylated RNAPII (RNAPIIo), which was confirmed by reciprocal pull-down of RNAPIIo (Fig. 1h, i). The RNAPIIo-ELOF1 interaction was constitutive and not affected by UV irradiation, demonstrating that, like its yeast counterpart, ELOF1 is an RNAPII-associated factor. The ELOF1^S72/D73K^ mutant failed to interact with RNAPIIo revealing that ELOF1 interacts with RNAPII in the same manner as its yeast ortholog ELF1. Importantly, stable expression of ELOF1^WT^ or ELOF1^ΔN^ fully rescued the Illudin S-sensitive phenotype of *ELOF1*-KO cells, whereas the ELOF1^S72/D73K^ mutant failed to restore this phenotype (Fig. 1j). These findings show that the interaction between RNAPII and ELOF1 is required to provide resistance against transcription-blocking DNA damage, while binding of ELOF1 to DNA through its N-terminal region is dispensable for this function.

### Loss of ELOF1 decreases transcription at the distal end of long genes without affecting RNAPII elongation rates

We next sought to gain insight into the role of ELOF1 under conditions without exogenous DNA damage. To this end, we performed a genome-wide CRISPR screen in *ELOF1*-KO RPE1-iCas9 cells to identify genes that cause synthetic lethality or synthetic viability in combination with knockout of *ELOF1*. This approach identified synthetic lethality (FDR<0.1) between *ELOF1* and the transcription elongation factors *SUPT4H1* (*SPT4* in yeast), *NELFE, INTS6* and *INTS12*, and the PAF1 subunits *CTR9* and *LEO1* (Fig. 2a, Fig S4a). At a Z-score cut-off of -3, related physically interacting proteins of the top hits were found, including *TFIIS (TCEA1)* and *SUPT6H* (Fig. 2b). This is in line with earlier work in yeast showing that ELF1 shows synthetic lethality with SPT4, PAF1 subunits, and TFIIS (Prather et al., 2005), likely because these elongation factors work together with yeast ELF1 to facilitate RNAPII progression through nucleosomes (Ehara et al., 2019; Ehara et al., 2017).

**Fig. 2.**
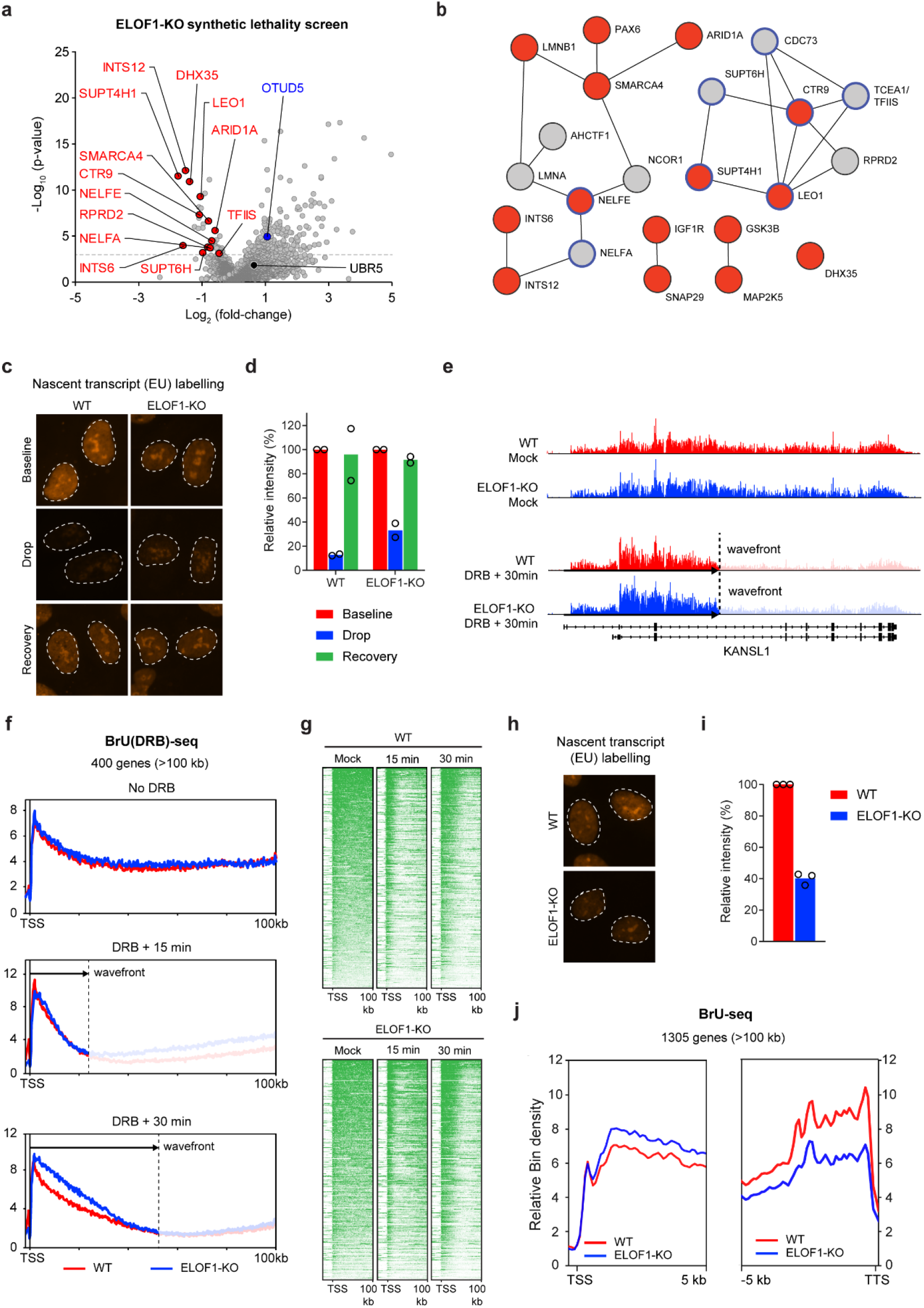
ELOF1 is required for processive transcription until the end of genes. (**a**) Volcano plot depicting gene-knockouts depleted (selected genes highlighted in red) or enriched (selected genes highlighted in blue) in proliferating *ELOF1*-KO cells as compared to WT control (x-axis, log_2_ fold-change), and their significance (y-axis, -log_10_ p-value; full analysis results in supplementary data). See Fig S4a for single guide counts. (**b**) Network of interacting hit genes (FDR<0.1, red) depleted in ELOF1-KO cells compared to WT cells, and depleted interactors (normalized Z- score<-3, gray).Blue edges reflect RNAPII interactors, gray lines indicate other protein-protein interactions. (**c**) Representative images of RPE1-iCas9 WT and *ELOF1*-KO cells after pulse-labelling for 1h with 5-EU to visualize nascent transcripts. Cells were either incubated with 5-EU (Baseline), incubated with DRB for 1h followed by incubation with DRB + 5-EU for 1h (Drop), or incubated with 5-EU after wash-out of DRB (Recovery). (**d**) Quantification of 5-EU levels normalized to the baseline level before DRB treatment for each condition. Each symbol represents the mean of an independent experiment (*n=2*), each containing 2 technical replicates, >80 cells collected per technical replicate. (**e**) UCSC genome browser track showing read density of BrU signal across the *KANSL1* gene after mock treatment, or at 30 m after DRB wash-out showing transcription wavefronts in WT (red) and *ELOF1*-KO cells (blue). (**f**) Metaplots of nascent transcription in 400 genes of >100 kb in WT (red) or *ELOF1*-KO (blue) cells after mock treatment, or after DRB wash-out (15 min or 30 min). (**g**) Heatmaps of BrU-seq data from the TSS into the first 100 kb of 400 genes of >100kb, ranked according to BrU signal in mock-treated cells (left panel). Heatmaps of the same genes are shown at 15 m or 30 m after DRB wash-out in WT or *ELOF1*-KO cells (middle and right panel). (**h**) Representative images of RPE1-iCas9 WT and *ELOF1*-KO cells after pulse-labelling for 1h with 5-EU to visualize nascent transcripts. (**i**) Quantification of 5-EU levels of *ELOF1*-KO cells normalized to WT cells shows that *ELOF1*-KO cells have reduced transcription levels compared to WT cells (*n=3*), each containing 2 technical replicates, >80 cells collected per technical replicate. (**j**) Metaplots of BrU signal (nascent transcription) in 1305 genes of >100 kb in WT (red) or *ELOF1*-KO (blue) cells in the first 5 kb after the TSS (left panel) and the last 5 kb before the TTS.

To investigate whether ELOF1 has a role in transcription elongation, we treated human cells with the reversible transcription inhibitor 5,6-dichlorobenzimidazole 1-beta-D ribofuranoside (DRB), and subsequently measured recovery of nascent transcription by 5-ethynyl-uridine (EU) labelling after wash-out of DRB (Fig. S4b). Although transcription was severely inhibited by treatment with DRB, both WT and *ELOF1*-KO cells showed a complete recovery of transcription within 1 h after removal of DRB (Fig. 2c, d). To more precisely analyse RNAPII elongation rates, we employed a variant of Bromouridine (BrU)-seq combined with a DRB pulse, hereafter called BrU(DRB)-seq. Briefly, cells were treated for 60 min with DRB followed by DRB wash-out to trigger the synchronized restart of transcription elongation into gene bodies. Following DRB removal, nascent transcripts were labelled with BrU for either 15 or 30 min and BrU-labelled RNA was isolated and subjected to deep sequencing. Transcription elongation rates were determined by measuring how far in the gene body transcription proceeded after removal of DRB (Veloso et al., 2014). An example of a representative gene, *KANSL1* (∼200 kb), is shown in Fig. 2e. Averaging the BrU(DRB)-seq profiles of 400 genes revealed that RNAPII progressed ∼25 kb into gene bodies at 15 min after DRB removal, and further to ∼50 kb after 30 min (Fig. 2f). This is consistent with the reported RNAPIIo elongation rate of ∼2 kb/min (Danko et al., 2013). Importantly, knockout of *ELOF1* did not affect RNAPII elongation rates after DRB wash-out at both time-points (Fig. 2f; see Fig 2e for the *KANSL1* gene). Heatmaps of BrU-labelled RNA from the TSS until 100 kb into the gene body of 400 selected genes of at least 100 kb indeed showed a very similar profile in wild-type and *ELOF1*-KO cells after DRB washout (Fig. 2g), indicating that knockout of ELOF1 does not affect transcription elongation in diploid human RPE1 cells. In line with this, *ELOF1*- KO cells did not display differences in chromatin accessibility at the TSS of ∼3000 genes measured by ATAC-seq (Fig. S4c).

Interestingly, when we quantified basal transcription based on 5-EU incorporation followed by microscopic analysis, we noted a ∼60% overall reduction in *ELOF1*-KO cells compared to wild-type cells (Fig. 2h, i; note that plots in Fig 2d are normalized to baseline for each cell-line). In line with this, analysis of nascent transcription by BrU-seq in 1305 genes of at least 100 kb revealed similar nascent transcript levels in the first 5 kb, while nascent transcription was considerably reduced in the last 5 kb of the same genes in *ELOF1*-KO cells (Fig. 2j, Fig S4d). Importantly, this effect was much milder for shorter genes and was hardly seen in genes between 25 and 50 kb (Fig S4d). Very similar to our findings on ELOF1, earlier work showed that depletion of SPT5 in mouse cells did not affect RNAPII elongation rates, but did cause reduced transcription in the distal ends of long genes (Fitz et al., 2018). We propose that, in the absence of ELOF1, RNAPII has difficulty progressing through inherent DNA structures, or possibly DNA lesions from endogenous sources.

### ELOF1 is required for transcription restart following genotoxic stress

DNA damage in actively transcribed strands causes a genome-wide transcriptional arrest due to RNAPIIo stalling at these lesions (Brueckner et al., 2007). A hallmark of TCR-deficient cells is the inability to resume transcription following UV irradiation due to a failure to remove transcription-blocking DNA lesions (Mayne and Lehmann, 1982).

To assess whether ELOF1 is involved in the restart of transcription, we measured the recovery of RNA synthesis (RRS) by 5-EU labelling at different time-points after UV irradiation. Microscopic analysis showed a strong transcriptional arrest to ∼30% residual nascent transcription at 3 h after UV (9 and 12 J/m^2^) in all cell lines (Fig. 3a, b, Fig. S5a). While WT cells completely recovered RNA synthesis within 24 h after UV irradiation, the TCR-deficient *CSB*-KO cells failed to restart transcription. Strikingly, *ELOF1*-KO cells showed a strong RNA synthesis recovery defect after UV irradiation, which was comparable to the defect in *CSB*-KO cells (Andrade-Lima et al., 2015) (Fig. 3a, b, Fig. S5a).

**Fig. 3.**
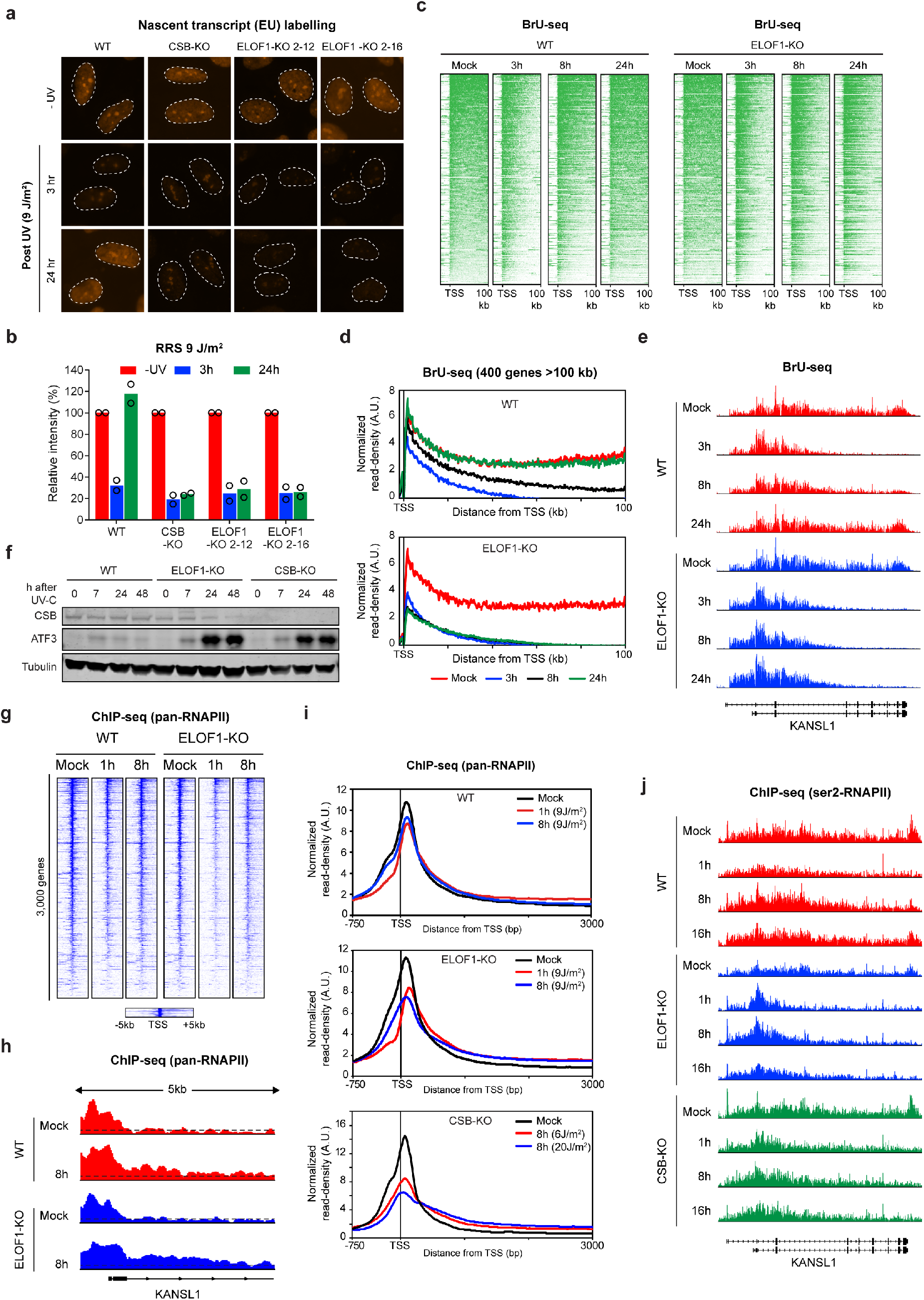
ELOF1 is essential for transcription recovery after UV. (**a**) Representative images of the indicated RPE1-iCas9 cells after pulse-labelling for 1 h with 5-EU to visualize nascent transcripts. Cells were either mock-treated (−UV) or UV-irradiated (3 h or 24 h; 9 J/m^2^). (**b**) Quantification of 5-EU levels normalized to mock-treated levels for each cell-line. Each symbol represents the mean of an independent experiment (*n=2*), each containing 2 technical replicates, >80 cells collected per technical replicate. (**c**) Heatmaps of BrU-seq data from the TSS into the first 100 kb of 400 genes of >100kb ranked according to BrU signal in mock-treated cells (left panel). Heatmaps of the same genes at 3 h, 8 h, and 24 h after UV irradiation (9 J/m^2^). (**d**) Metaplots of BrU signal (nascent transcription) in 400 genes of >100 kb in WT (upper) or *ELOF1*-KO (lower) cells after mock treatment (red), or 3 h (blue), 8 h (black), and 24 h (green) after UV irradiation (9 J/m^2^). (**e**) UCSC genome browser track showing read density of BrU signal across the *KANSL1* gene after mock treatment, or at 3h, 8 h, or 24 h after UV irradiation (9 J/m^2^) in WT (red) or *ELOF1*-KO cells (blue). (**f**) Western blot analysis of ATF3 protein levels in the indicated RPE1-iCas9 cell-lines after mock treatment, or 7 h, 24 h, and 48 h after UV irradiation (9 J/m^2^) (*n=4* for WT and *ELOF1*-KO, and n=3 for *CSB*-KO). (**g**) Heatmaps of pan-RNAPII ChIP-seq data around the TSS of 3,000 genes of 3-100kb, ranked according to RNAPII signal in mock-treated WT cells. Heatmaps of the same genes are shown at 1 h or 8 h after UV irradiation (9 J/m^2^) in WT or *ELOF1*- KO cells. (**h**) UCSC genome browser track showing read density of pan-RNAPII signal in the first 5kb of the *KANSL1* gene after mock treatment, or at 8h after UV irradiation (9 J/m^2^) in WT (red) and *ELOF1*-KO cells (blue). (**i**) Averaged metaplots of pan-RNAPII ChIP-seq of 3,000 genes of 3-100 kb around the TSS in RPE1-iCas9 WT (upper) or RPE1 *ELOF1*-KO (middle) after mock- treatment (black) or at 1 h (red) or 8 h (blue) after UV irradiation (9 J/m^2^). The lower panel show metaplots in U2OS *CSB*-KO cells after mock-treatment (black) or at 8 h after UV irradiation with either 6 J/m^2^ (red) or 20 J/m^2^ (blue). (**j**) UCSC genome browser track showing the read density of the ser2-RNAPII signal across the *KANSL1* gene after mock treatment, or at 1 h, 8 h or 16 h after UV irradiation (9 J/m^2^) in WT (red), *ELOF1*-KO (blue) or *CSB*-KO cells (green).

To further explore the recovery of RNA synthesis in a genome-wide manner, we labelled nascent transcripts with BrU at multiple timepoints after UV irradiation (3, 8, 24 hrs) followed by isolating and sequencing of BrU-labelled RNA. Nascent transcription was strongly reduced 3 h after UV irradiation at the TSS and decreased progressively toward more distal genic regions, with a virtual loss of transcription beyond 100 kb into gene bodies, in both WT and *ELOF1*-KO cells (Fig. 3c, d). Although milder, a similar loss of nascent transcription was also observed toward the end of shorter genes (25-50 kb and 50-100 kb; Fig. S5b), which fits with the distribution of transcription-blocking photolesions (Perdiz et al., 2000), and the probability that RNAPII encounters such a lesion. While nascent transcription was partially resumed 8 h after UV irradiation and fully restored 24 h after UV irradiation in WT cells, *ELOF1*-KO cells failed to recover (Fig. 3c, d, see Fig. 3e f or the representative *KANSL1* gene). These findings demonstrate that ELOF1 has an essential role in transcription recovery following genotoxic stress.

### Reduced binding of RNAPII at promoters and increased stalling in *ELOF1*-KO cells after UV

Genotoxic stress induces the expression of the transcription repressor ATF3, which downregulates the expression of ∼ 3000 genes harboring a CRE/ATF binding site (Epanchintsev et al., 2017). Both CSB and CSA promote the degradation of ATF3 thereby relieving its inhibitory impact on transcription initiation and enabling transcription restart (Epanchintsev et al., 2017). Western blot analysis revealed an increase in the steady-state ATF3 protein levels 7 h after UV irradiation in WT, *ELOF1*-KO and *CSB*-KO cells (Fig. 3f). The ATF3 protein levels decreased in WT cells within 48 h after UV irradiation, while ATF3 protein levels increased dramatically in both *ELOF1*- KO and *CSB*-KO cells at 24 h and 48 h after UV (Fig. 3f). These findings may provide an explanation for the reduced RNA synthesis at TSS sites following UV irradiation in *ELOF1*-KO cells (Fig. 3d, S5b). Interestingly, RNA-sequencing (RNA-seq) showed that ATF3 transcript levels were strongly upregulated 24 h after UV irradiation in *ELOF1*-KO and *CSB*-KO cells compared to WT cells (Fig. S6a-c). This suggests that increased ATF3 expression at late time-points after UV in these cells is at least in part controlled at the mRNA synthesis level. In addition, while *ELOF1* and *CSB* knockout cells showed a strong upregulation of short pro-survival genes, such as *CDKN1A* (Bugai et al., 2019), a set of longer genes was downregulated at 24 h after UV (Fig S6a-d), consistent with the RNA synthesis recovery defect in these cells (Fig. 3a, b).

To monitor transcription initiation more directly, we combined chromatin immunoprecipitation with sequencing (ChIP-seq) using antibodies against unmodified RNAPII (pan-RNAPII) to map genome-wide binding of RNAPII following UV irradiation. Heatmaps around the TSS showed a strong signal for ∼3000 genes around the promoter in non-irradiated cells, which was comparable between wild-type and *ELOF1*-KO cells (Fig. 3g). At 1 h and 8 h after UV irradiation, we detected a striking reduction in RNAPII binding at TSS sites in *ELOF1*-KO cells, but not in WT cells (Fig. 3g, h, i), which was accompanied by an increase in RNAPII reads in the gene body (Fig. 3i, Fig S5c). A highly similar redistribution of RNAPII after UV irradiation was detected in *CSB*-KO cells, suggesting that this corresponds to increased stalling at DNA lesions in gene bodies (Fig. 3i). This is consistent with our BrU-seq data showing that RNA synthesis is already largely restored around TSS sites at 8 h after UV in WT cells, but not in *ELOF1*-KO cells (Fig. 3c, d).

We also detected enrichment of pan-RNAPII after transcription termination sites (TTS) in non-irradiated cells, reflecting post-transcriptional pausing prior to dissociation (Fig. S5c, d). However, this enrichment was strongly reduced in *ELOF1*-KO cells at 8 h after UV irradiation, but not in WT cells, which is consistent with fewer transcripts reaching the end of genes (Fig. S5c, d). In line with this, ChIP-seq experiments using antibodies against elongating RNAPII (Ser2- RNAPII), revealed that reads throughout gene-bodies and at TTS sites indeed recovered within 16 h after UV in WT cells, but not in *ELOF1*-KO and *CSB*-KO cells (Fig. 3j). These findings suggest that ELOF1 is either directly involved in the repair of UV-induced DNA lesions or in the restart of transcription after these lesions are repaired by TCR.

### The genetic network of ELOF1 and CSB in response to DNA damage

To further compare the roles of CSB and ELOF1 in the response to Illudin S-induced DNA damage and their connections to known TCR factors, we repeated the Illudin S screen in *CSB*-KO and *ELOF1*-KO cells at reduced Illudin S concentrations (2 nM for *CSB*-KO and 4 nM for *ELOF1*-KO). Comparing the drug-genetic interactions in *CSB*-KO cells to those of wild-type cells after Illudin S exposure revealed that, as expected, *CSA* and *UVSSA* loss did not cause an additive sensitivity to Illudin S (FDR>0.9). Interestingly, loss of *ELOF1*, as well as the known DNA damage-induced transcription regulators *DOT1L* (Oksenych et al., 2013) and *HIRA (Adam et al*., *2013)*, did cause additive sensitivity to Illudin S (FDR<0.001, Fig. 4a, S5e). The screen in *ELOF1*-KO cells confirmed the additive sensitivity by loss of the *CSA, CSB* or *UVSSA* core TCR genes (FDR<0.01), while loss of *DOT1L* and *HIRA* did not further sensitize *ELOF1*-KO cells (FDR>0.9, Fig. 4b, S5e), suggesting that ELOF1, in addition to TCR, also acts in a second compensatory pathway, possibly together with *HIRA* and *DOT1L*. Interestingly, loss of STK19, which was previously linked to transcription recovery after UV (Boeing et al., 2016) and strongly sensitizes RPE1 cells to Illudin (Fig. 1b,c), did not cause additional Illudin S sensitivity in either *CSB*-KO or *ELOF1*-KO cells (FDR>0.9 and FDR>0.5 respectively), indicating that STK19 acts downstream of both CSB and ELOF1 (Fig. 4a, b). Our analysis places all the currently known transcription recovery genes in a genetic framework (Fig. 4c). Interestingly, both *CSB*-KO and *ELOF1*-KO cells revealed strong additive Illudin S sensitivity upon loss of REV7 or the 9-1-1 complex (Fig. 4a, b), suggesting that these KO cells are more dependent on translesion synthesis to deal with unresolved transcription-blocking DNA damage acquired during replication (Fig. 4c). To validate these genetic interactions, we generated *CSB/ELOF1* and *CSB/CSA* double knockout (dKO) cells. Drug-sensitivity assays confirmed that *CSB/ELOF1*-dKO cells were more sensitive to Illudin S and Irofulven than either single knockout, while this was not the case for *CSB/CSA*-dKO cells (Fig. 4d, e). The accompanying paper by Geijer *et al*. shows that yeast ELF1, similar to human ELOF1, is involved in both RAD26-dependent and RAD26-independent repair pathways, suggesting that this dual role is conserved throughout evolution.

**Fig. 4.**
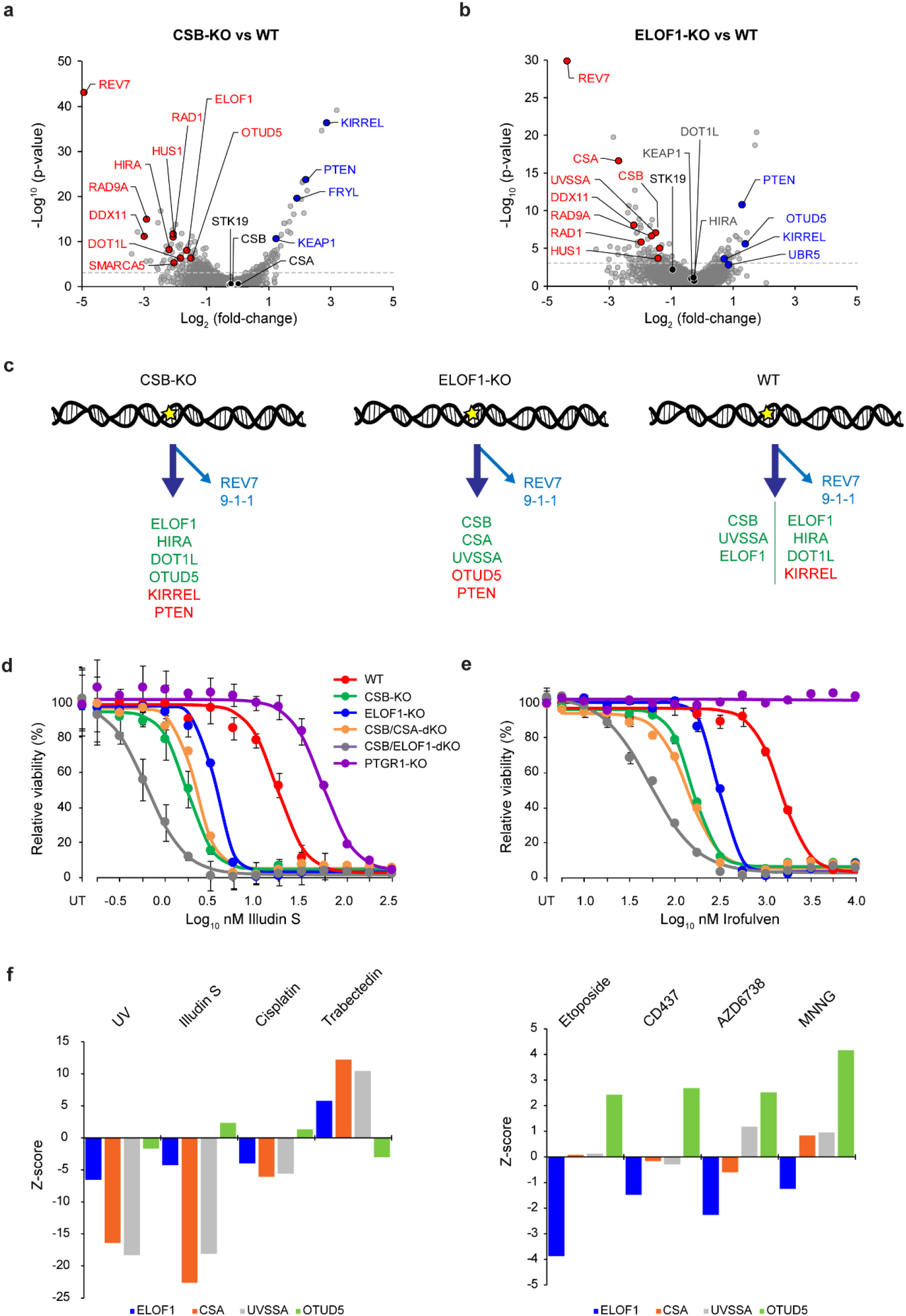
CRISPR screens identify determinants of Illudin S sensitivity in the absence of ELOF1 or CSB. (**a, b**) Volcano plot depicting gene-knockouts sensitizing (selected genes highlighted in red) or conferring resistance (selected genes highlighted in blue) Illudin S treatment (x-axis, log_2_ fold-change), and their significance in the *ELOF1*-KO (a) and CSB-KO (b) cell lines.(y-axis, - log_10_ p-value; full analysis results in Supplementary data). (**c**) Scheme depicting the identified DNA repair pathways. (**d, e**) 72 h drug-sensitivity assaysin the indicated RPE1-iCas9 single and double KO clones exposed to (**d**) Illudin S (*n=3*) or (**e**) Irofulven (*n=3*). Values indicate the average ± SEM of 4 technical replicates from one experiment. (**f**) Results of mining a recent CRISPR screen repository. Shown are the Z-scores for the indicated sgRNAs (targeting *ELOF1* (blue), *CSA* (orange), *UVSSA* (grey), *OTUD5* (green) after exposure to the indicated genotoxic agents.

Interestingly, our screens reveal that loss of the *PTEN* tumor suppressor strongly reduces the cytotoxic effects of transcriptional stress in both *ELOF1*-KO and *CSB*-KO cells (Fig. 4a-c). Furthermore, our genetic-interaction maps also show that knocking out the deubiquitylase *OTUD5* alleviates the sensitivity of *ELOF1*-KO cells to Illudin S, while its loss has the opposite impact on *CSB*-KO cells (Fig. 4a, b, S5e). Linking this enzyme to the transcription-stress response, OTUD5 was recently found to repress RNAPII elongation in response to DNA double-strand breaks (de Vivo et al., 2019). By mining a recently published resource involving genome-wide CRISPR screens in response to 25 genotoxic agents (Olivieri et al., 2019), we found that *ELOF1* loss indeed showed a similar response as *CSA* and *UVSSA* loss to compounds that induce transcription-blocking DNA lesions, such as Illudin S (Jaspers et al., 2002), UV light, Cisplatin (Enoiu et al., 2012), and Trabectedin (Takebayashi et al., 2001) (Fig. 4f). However, knockout of *ELOF1* caused a profound sensitivity to DNA lesions that provoke a DNA replication-stress response, such as Etoposide, ATR inhibitor, and DNA polymerase α inhibitor. This sensitivity is not observed in *CSA* and *UVSSA* knockouts (Fig. 4f). Strikingly, *OTUD5* loss often showed the opposite impact to *ELOF1* loss in response to these genotoxic agents (Fig. 4f). These findings indicate that ELOF1, in addition to its role in canonical TCR, also acts in an OTUD5-regulated parallel TCR pathway together with HIRA and DOT1L (Fig. 4c), for instance by preventing collisions between transcription and replication machineries. Thus, ELOF1 appears to be a general sensor of transcriptional stress.

### Genome-wide TCR-seq reveals that ELOF1 is an essential TCR factor

The strong RNA synthesis defect in *ELOF1*-KO cells after UV irradiation is comparable to the impact of *CSB* loss. This can be explained by a role of ELOF1 in TCR-mediated DNA repair, or by a specific role in transcription restart after repair.

To distinguish between these possibilities, we measured genome-wide TCR kinetics using a strand-specific ChIP-seq technology we recently developed (TCR-seq) (Nakazawa et al., 2020). To this end, we performed ChIP-seq experiments at different time-points after UV (1, 4, 8, 16 hrs) using an antibody recognizing RNAPIIo. The presence of DNA lesions in the transcribed strand causes stalling of RNAPIIo. The DNA fragments that are co-purified with RNAPIIo after ChIP are therefore highly enriched for UV-induced photolesions specifically in the transcribed (template) strand, but not in the coding strand (Brueckner et al., 2007). During NGS library preparation after ChIP, we ligated asymmetrical adapters to preserve strand-specific information. Importantly, UV-induced photolesions block PCR amplification during library preparation resulting in a loss of reads in the template (transcribed) strands in the presence of DNA lesions (Fig. 5a) (Nakazawa et al., 2020). Thus, TCR-seq allows the strand-specific PCR amplification of fragments without DNA damage, enabling a genome-wide quantification of TCR kinetics.

**Fig. 5.**
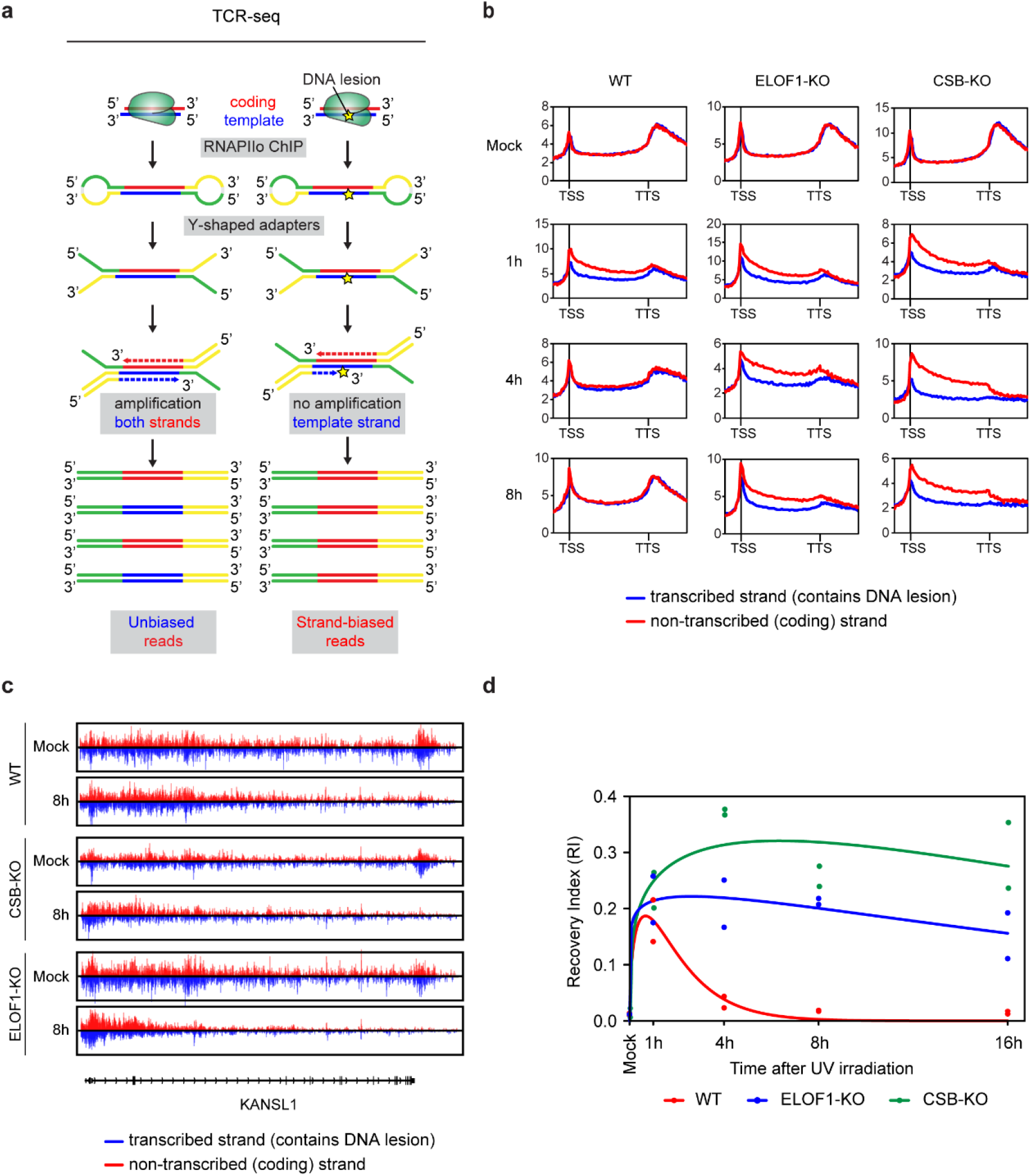
ELOF1 is essential for repair of transcription-blocking DNA damage. (**a**) Schematic representation of TCR-seq. (**b**) Averaged metaplots of Ser2-RNAPII TCR-seq of 3,000 genes from the TSS until the TTS (−5kb, +5kb respectively) in the indicated RPE1-iCas9 cells after mock-treatment or at 1h, 4h, or 8h after UV irradiation (9 J/m^2^). The coding (non-transcribed) strand is shown in red, while the template (transcribed) strand is shown in blue (*n=2*). See Fig S7a for individual replicates and additional timepoints. (**c**) UCSC genome browser track showing read densities of TCR-seq data based on strand-specific Ser2-RNAPII signal across the *KANSL1* gene after mock treatment, or at 8 h after UV irradiation (9 J/m2) in WT, *CSB*-KO or *ELOF1*-KO cells. Reads from the coding (non-transcribed) strand are shown in red, while reads from the template (transcribed) strand are shown in blue. See Fig S7b for additional time points. (**d**) Time-course of the indicated RPE1-iCas9 cells showing the recovery index (RI), representing genome-wide TCR repair kinetics. Per sample, a frequency distribution plot was generated of the per-gene strand specificity index (SSI; defined as the relative difference in read density between the transcribed and non-transcribed strands) of 3,000 genes of 3-100kb. The RI is subsequently obtained by fitting a mixture of 3 Gaussian distributions, corresponding to undamaged gene fractions (SSI=0) and two unrepaired gene fractions (i.e. |SSI|<>0) (see Fig S8a,b). The RI was calculated from duplicate time-course experiments shown in Fig S8a.

TCR-seq revealed a strong strand-bias in WT, *ELOF1*-KO, and *CSB*-KO cells 1 h after UV irradiation which was not present in non-irradiated cells (Fig. 5b, Fig. S7a, see *KANSL1* gene in Fig. 5c, Fig. S7b). Strikingly, while this strand-bias was fully resolved in wild-type cells over the course of 16 h, both *ELOF1*-KO and *CSB*-KO cells still displayed a strong strand-bias at all time-points analyzed (Fig. 5b,c Fig. S7a,b, Fig S8a). Resolving the strand-bias in WT cells was accompanied by the reappearance of Ser2-RNAPII reads throughout gene bodies and after the TTS site, which did not occur in the absence of *ELOF1* or *CSB* (Fig. 5b, c, Fig. S7a, b). Analysis of the recovery index (RI) (based on data shown in Fig S8a, see also Fig S8b), which is a measure for the genome-wide removal of DNA lesions from transcribed strands showed that WT cells cleared transcription-blocking DNA lesions from the genome within 16 h, while *ELOF1*-KO and *CSB*-KO cells displayed no significant repair within this timeframe (Fig. 5d). Strand-specific analysis after ChIP-seq with another (pan-RNAPII) antibody confirmed these results (Fig. S9a, b). These findings show that RNAPII persistently stalls at DNA lesions in *ELOF1*-KO cells, and identify ELOF1 as a new core factor in TCR. To address if ELOF1 is also involved in global genome repair (GGR), we measured unscheduled DNA synthesis (UDS) following global UV irradiation. Importantly, *ELOF1*-KO cells were fully capable of removing lesions throughout the genome (Fig. S10a, b), demonstrating that ELOF1 is specifically involved in TCR.

### TCR complex assembly and RNAPII ubiquitylation

Having identified ELOF1 as a new core repair factor in TCR, we next investigated whether ELOF1 is involved in TCR complex assembly. To this end, we immunoprecipitated endogenous RNAPIIo by employing a procedure we recently established to isolate intact TCR complexes (van der Weegen et al., 2020). Both CSB and CSA associated with RNAPII in a UV-specific manner in wild-type as well as *ELOF1-KO* cells, while the association of CSA with RNAPIIo was completely abolished in cells lacking *CSB* (Fig. 6a). The latter was expected since CSB is the first protein to be recruited to damage-stalled RNAPIIo (van der Weegen et al., 2020). Although TFIIH (p89 and p62 subunits) associated in a UV-specific manner with RNAPIIo in wild-type cells, the interaction was severely reduced in *ELOF1-KO* cells in both RPE1 and U2OS cells (Fig. 6a, b). Importantly, re-expressing ELOF1-GFP in *ELOF1-KO* cells completely restored TFIIH recruitment to DNA damage-stalled RNAPIIo (Fig. 6b).

**Fig. 6.**
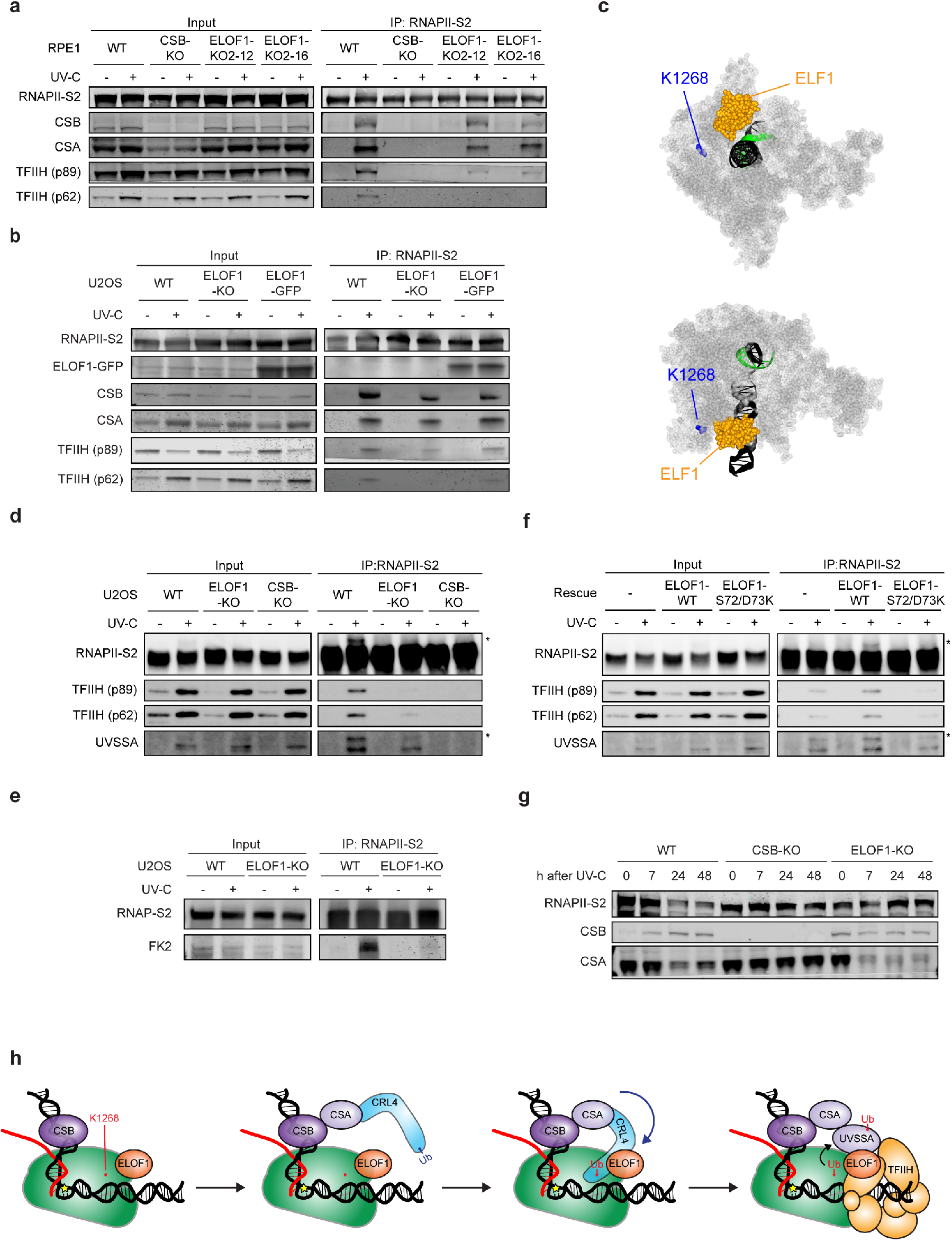
ELOF1 directs RNAPII ubiquitylation. (**a**) Endogenous RNAPIIo Co-IP on RPE1iCas9 WT, *CSB*-KO, and *ELOF1*-KO clones (*n=3)*. (**b**) Endogenous RNAPIIo Co-IP on U2OS (FRT) WT, *ELOF1*-KO and *ELOF1*-KO complemented with ELOF1^WT^-GFP (*n=2*). (**c**) Front-view and top-view of the structure (PDB: 5XON) of *K. pastoralis* ELF1 (orange) bound to RNAPII (grey). The RPB1-K1286 ubiquitylation site (K1264 in *K. pastoralis*) is indicated in blue. (**d**) Endogenous RNAPIIo Co-IP on U2OS (FRT) WT, *ELOF1*-KO, and *CSB*-KO clones. The ubiquitylated form of RNAPII and UVSSA, detected as higher migrating bands, are indicated with an asterisk. (**e**) Endogenous RNAPIIo Co-IP on U2OS (FRT) cells (WT and *ELOF1*-KO) in the presence of deubiquitylase inhibitor N-ethylmaleimide (NEM). The ubiquitylated form of RNAPII is detected using FK2 antibody. (**f**) Endogenous RNAPIIo Co-IP on U2OS (FRT) *ELOF1*-KO and *ELOF1*-KO complemented with ELOF1^WT^-GFP or ELOF1^S72/D73K^-GFP. (**g**) Western blot analysis of CSA protein levels of the indicated RPE1-iCas9 cell-lines after mock treatment, or 7 h, 24 h, and 48 h after UV irradiation (9 J/m^2^). (**h**) Model of how ELOF1 serves as a specificity factor for RNAPII ubiquitylation during TCR.

We have recently shown that RNAPII is ubiquitylated on lysine K1268 of the RPB1 subunit in response to UV irradiation and that the CRL^CSA^ complex is the key E3 ubiquitin ligase required for this modification. Moreover, RPB1-K1268 ubiquitylation is required to recruit the TFIIH complex (Nakazawa et al., 2020). It is interesting to note that the yeast ortholog ELF1 is bound to RNAPII in close proximity to the K1268 (K1264 in *K. pastoralis*) ubiquitylation site (Fig. 6c, Fig S1c). We therefore hypothesized that ELOF1 may regulate the UV-induced ubiquitylation of RNAPII. While immunoprecipitation of endogenous RNAPIIo showed that RNAPII is ubiquitylated in response to UV in wild-type cells, this UV-induced modification is absent in cells lacking either *ELOF1* or *CSB* (Fig. 6d). Immunoprecipitation of endogenous RNAPIIo following detection of endogenous ubiquitin conjugates using an FK2 antibody (Fig. 6e), or by detection of Flag-tagged ubiquitin confirmed this phenotype (Fig. S10c).

While endogenous UVSSA associated with DNA damage-stalled RNAPIIo in WT cells, its recruitment was decreased in *ELOF1-KO* cells, and virtually absent in *CSB-KO* cells. Interestingly, although residual UVSSA association with RNAPII was detected in *ELOF1-KO* cells, we noticed that the upper band, corresponding to monoubiquitylated *UVSSA* (Nakazawa et al., 2020), was strongly reduced in *ELOF1*-KO cells (Fig. 6d). This is in line with previous work demonstrating that RNAPII ubiquitylation supports the ubiquitylation of UVSSA and that, in turn, UVSSA ubiquitylation is required for TFIIH recruitment (Nakazawa et al., 2020). Importantly, the loss of UV-induced RNAPII ubiquitylation in *ELOF1*-KO cells was fully restored by the stable expression of ELOF1^WT^, while expression of ELOF1^S72/D73K^ failed to rescue RNAPII ubiquitylation, UVSSA mono-ubiquitylation, as well as the association of TFIIH with DNA damage-stalled RNAPIIo (Fig. 6f). This shows that the interaction between RNAPII and ELOF1 is required during all of these steps. In line with a role for ELOF1 in RNAPII ubiquitylation and subsequently recruitment of UVSSA, western blot analysis revealed that CSA is almost completely degraded at later time-points (7, 24, 48 hrs) after UV irradiation in *ELOF1-KO* cells but not in WT or *CSB-KO* cells (Fig. 6g). This suggests that ELOF1 prevents proteasomal degradation of CSA through an auto-ubiquitylation process. A similar CSA degradation phenotype was also observed in *UVSSA*-KO cells (Fig. S10d), suggesting that CSA is de-ubiquitylated by the UVSSA-binding partner USP7, which is defective in *ELOF1*-KO cells as a result of the decreased recruitment of ubiquitylated UVSSA.

Together, our data indicate that ELOF1 acts as an RNAPII-associated specificity factor that directs the ubiquitin ligase activity of the CRL^CSA^ complex to facilitate ubiquitylation of K1268 on RNAPII, close to the ELOF1-binding site. This, in turn, promotes the recruitment and ubiquitylation of UVSSA in order to transfer the TFIIH complex onto DNA damage-stalled RNAPII to initiate repair (Fig. 6h).

## Discussion

The yeast orthologue of human ELOF1, called ELF1, stimulates RNAPII progression through nucleosomes together with elongation factors SPT4 and SPT5 (SUPT4H1 and SUPT5H in humans) in a manner depending on TFIIS to reactivate stalled RNAPII (Ehara et al., 2019). The impact of ELF1 on RNAPII progression *in vitro* is very mild, but there is strong cooperativity between ELF1, SPT4/SPT5 and TFIIS (Ehara et al., 2019). Through genetic-interaction mapping by isogenic CRISPR screens, we find that in the absence of genotoxic stress, ELOF1 knockout displays synthetic effects with loss of multiple transcription elongation factors and their interactors, comprising an interaction network including SUPT4H1, SUPT6H, subunits of the PAF1, NELF, and INTS complexes, and TFIIS (Fig. 2a, S4a). These findings suggest that a similar cooperativity exists in human cells. Of interest, the extended C-terminal acidic tail of yeast ELF1, which binds to histones H2A-H2B and may be required for nucleosome reassembly after RNAPII progression through nucleosomes (Ehara et al., 2019), is absent in human ELOF1, suggesting that this role is not conserved throughout evolution. Similarly, the basic N-terminal region, which may weaken histone-DNA contacts and facilitate transcription through nucleosomes in yeast (Ehara et al., 2019), is not required for the repair function of human ELOF1 (Fig. 1j).

The rate of RNAPII elongation was not affected in *ELOF1*-KO cells, likely due to functional redundancy with other elongation factors (Fig. 2e-g), similar to what has been observed for SUPT5H in mammalian cells (Fitz et al., 2018). However, loss of ELOF1 did specifically reduce transcription in the distal ends of genes without affecting initiation (Fig. 2j, S4d), suggesting that RNAPII molecules need ELOF1 to reach the end of longer genes. We favor the scenario that RNAPII has difficulty progressing through inherent DNA structures or DNA lesions caused by unknown endogenous sources resulting in reduced transcription towards the end of longer genes (Fig. 2a, b).

Structural studies have revealed that yeast ELF1 binds to a central cleft in front of RNAPII where the DNA enters the enzyme (Ehara et al., 2017) (Fig. 6c. Fig S1b, c). This binding site is very close to a recently identified UV-induced ubiquitylation site in RNAPII (RPB1-K1268), which is essential for the efficient repair of and recovery from transcription-blocking DNA lesions (Nakazawa et al., 2020; Tufegdžić Vidaković et al., 2020). Our experiments show that the interaction between RNAPII and ELOF1 is indeed essential for the UV-induced ubiquitylation of DNA damage-stalled RNAPII (Fig. 6d-f). Together, our findings suggest a model in which ELOF1 constitutively interacts with elongating RNAPII, and promotes its reactivation to ensure progression past obstacles that RNAPII can overcome without TCR. The stalling of RNAPII at DNA lesions that block transcription and depend on TCR for repair trigger the sequential recruitment of CSB and CSA, as well as the CSA-associated CRL complex (CRL^CSA^). Loss of ELOF1 does not affect the recruitment of either CSB or CRL^CSA^ to lesion-stalled RNAPII, but we propose that ELOF1 directs the catalytic site of the CRL^CSA^ into close proximity of the K1268 site. By serving as a specificity factor for RNAPII ubiquitylation, ELOF1 subsequently promotes the recruitment and ubiquitylation of UVSSA, which, in turn, mediates the transfer of the TFIIH complex from UVSSA onto DNA damage-stalled RNAPIIo to initiate repair (Fig. 6h).

The importance of ELOF1 in TCR is evident from *(1)* the inability of *ELOF1*-KO cells to remove UV-induced lesions from transcribed strands (Fig. 5, Fig. S7 - Fig. S9), *(2)* reduced binding of RNAPII at promoters, but increased retention in gene bodies, following UV irradiation in *ELOF1*-KO cells, which is consistent with persistent stalling of RNAPII throughout the genome (Fig. 3h-j, Fig. S5c), and *(3)* the inability of *ELOF1*-KO cells to recover RNA synthesis following UV irradiation to a similar extent as TCR-deficient *CSB*-KO cells (Fig. 3a-d, Fig. S5a, b) (Mayne and Lehmann, 1982). We therefore propose that ELOF1 should be considered as a new core TCR factor, and we predict that inactivating mutations affecting ELOF1 function, if viable, would likely cause a Cockayne syndrome-like phenotype.

Although *ELOF1*-KO cells are already highly sensitive to transcription-blocking DNA damage, knockout of the other TCR core genes (*CSB, CSA* and *UVSSA*) sensitized *ELOF1*-KO cells even further, while loss of either *CSA* or *UVSSA* in *CSB*-KO cells did not cause additive sensitivity to Illudin S or Irofulven (Fig. 4a-e). These genetic-interactions indicate that ELOF1 is not only involved in TCR, but also in another compensatory pathway that deals with transcription stress. Like ELOF1, the chromatin modifiers HIRA and DOT1L, which play a role in epigenetic modifications in response to DNA damage (Adam et al., 2013; Oksenych et al., 2013), also become essential in Illudin S-treated *CSB* knock-out cells (Fig. 4a-c, S5e). This suggests that HIRA and DOT1L act in the same ELOF1-dependent compensatory pathway. Unlike CSB, these three genes provide protection against genotoxic agents causing replication stress (Olivieri et al., 2019) (Fig. 4f), suggesting that ELOF1 together with DOT1L and HIRA may act in a CSB-independent pathway that prevents collisions between transcription and replication machineries.

In yeast, two redundant TCR pathways are known, with one being dependent on RAD26 (the orthologue of CSB), and the other being dependent on the non-essential RNAPII subunit RPB9 (Li and Smerdon, 2002), which promotes RPB1 ubiquitylation in response to UV irradiation (Chen et al., 2007). Reactivation of TCR in RAD26-deficient yeast cells occurs when elongation factors SPT4 or SPT5 are also deleted (Ding et al., 2010; Jansen et al., 2000), suggesting that RAD26 in yeast may not be a TCR factor that recruits the repair machinery to stalled RNAPII. In contrast, CSB is essential to recruit the repair machinery, and plays an important role in RNAPII ubiquitylation in human cells through facilitating the recruitment of the CRL^CSA^ complex (Nakazawa et al., 2020; van der Weegen et al., 2020). Moreover, our genetic screen reveals that knockout of *SUPT5H* does not alleviate the sensitivity of *CSB*-KO cells, and that *POLR2I* (human orthologue of RPB9) knockout is important for viability, but does not cause Illudin S sensitivity in human cells (Fig. S10e). Interestingly, yeast RPB9 is bound to the RNAPII complex close to the K1268 (K1264 in *K. pastoralis*) ubiquitylation site where ELF1 is also located, suggesting that functional crosstalk is possible between these two RNAPII components (Fig. S1d). Despite these differences between yeast and human cells, we favor the scenario that another TCR pathway may exist in human cells and that ELOF1, in addition to its role in CSB-dependent TCR, also plays a role in this compensating TCR pathway. In line with this, *ELOF1*-KO cells are sensitive to a range of genotoxic compounds to which core TCR knockouts (*CSA, CSB, UVSSA*) show no sensitivity (Olivieri et al., 2019) (Fig 4f). This compensatory TCR pathway seems to be modulated by de deubiquitylase OTUD5, considering that knockout of this enzyme alleviated the Illudin S-sensitive phenotype of *ELOF1*-KO cells, while it caused additive sensitivity in *CSB*-KO cells (Fig 4a, b, S5e). One possibility is that OTUD5 acts upon ubiquitylated RNAPII (on K1268) and that remaining very low levels of this UV-induced mark in *ELOF1*-KO cells are stabilized when the opposing activity of OTUD5 is also lost. Alternatively, the reported role of OTUD5 in arresting RNAPII at DNA double-strand breaks by inhibiting the FACT chaperone may be relevant here (de Vivo et al., 2019), particularly considering that ELF1 in yeast cooperates with FACT during transcription elongation (Ehara et al., 2019). Elucidating the components and the mechanism involved in the compensatory ELOF1-dependent pathway, and the genes that influence cytotoxicity in response to transcriptional stress, such as *PTEN*, are important goals for future research.

Together, our results identify ELOF1 as a new transcription-coupled DNA repair factor that promotes CSB-dependent TCR as a specificity factor for RNAPII ubiquitylation and, at the same time, acts in a compensatory second pathway as a more general sensor of transcription stress.

## Acknowledgments

The authors acknowledge Angela Kragten, Marta San Martin Alonso, Sylvie Noordermeer, Kana Kato, Mayuko Shimada, Susan Kloet, Shivani Rampersad, Maja Vukic, Daniel Warmerdam, Job de Lange, Anisha Ramadhin and Lucia Daxinger for help during this project. We also thank Jason Moffat, Katie Chan and Amy Tong for sharing the pLCKO-TKOv3 library prior to publication and. We thank Josephine Dorsman for bio-informatics support and the Amsterdam UMC NGS sequencing facilities for sample processing and NGS data mapping.

## Funding

This work was funded by an LUMC Research Fellowship and an NWO-VIDI grant (ALW.016.161.320) to MSL, a Leiden University Fund (LUF) grant to DvdH (W18355-2-EM), a KWF/Alpe Young lnvestigator Grant 10701 grant (JvS), a CCA proof-of-concept grant (KdL, RW), an Amsterdam UMC Innovation Grant (CRISPR Expertise Center, 2019; RW), and UM1 HG009382 and R01 CA213214 NCI grants to ML.

## Author contributions

YvdW generated U2OS and RPE1-iCas9 single KO and double KO cells, validated these all KO cells, generated FlpIn and Flag-Ub cell-lines, performed clonogenic survivals, co-IP experiments in RPE1-iCas9 and U2OS, RRS and DRB-RRS experiments, western blot analyses, generated samples for pan-RNAPII ChIP-seq, ser2-RNAPII ChIP-seq, ATAC-seq, DRB-BrU-seq, and BrU-seq, and wrote the paper. KdL validated single cell Cas9 induction in RPE1-iCas9, optimized and performed all Illudin S and isogenic CRISPR screens and their statistical analysis, performed and analyzed UV/Mock RNA-seq experiments and drug-sensitivity assays in RPE1-iCas9 single KO, double KO cells, generated samples for the BrU-seq experiment, performed network analyses and wrote the paper. DvdH performed Co-IP experiments in RPE1, developed tools and analyzed pan-RNAPII ChIP-seq, ser2-RNAPII ChIP-seq, ATAC-seq, DRB-BrU-seq, and BrU-seq. YN performed Co-IP experiments. IVN analysed DRB-BrU-seq and BrU-seq. NK generated U2OS single KO clones, generated Flp-In cell-lines, performed clonogenic survivals, and Co-IP experiments in U2OS. APW performed UDS experiments. KR helped with the Illudin S ELOF1 and CSB screens, RNAseq data analysis and network mapping; JJMvS created RPE1-iCas9 cell line and helped with the Illudin S and ELOF1 essentialome screens. YH analyzed ser2-RNAPII ChIP-seq. ML supervised IVN, and analysed DRB-BrU-seq and BrU-seq. TO supervised YN and YH, and analysed Ser2-RNAPII ChIP-seq. RMFW supervised KdL, KR and JJMvS, conceived, coordinated and supervised the CRISPR screens and hit analyses and reviewed other parts of the project, and wrote the paper. MSL supervised YvdW, DvdH, NK, APW, conceived, coordinated, and supervised the project, and wrote the paper.

## Competing interests

Authors declare no competing interests.

## Data and materials availability

Both raw and processed ChIP-seq, BrU-seq, ATAC-seq, and RNA-seq data are deposited in the Gene Expression Omnibus (GEO) under GSE149760 (password: obmhawiqvnenxal). Additional data and custom code will be made available upon reasonable request.

## Notes

### Competing Interest Statement

The authors have declared no competing interest.

