## Supplementary Materials for "ELOF1 is a transcription-coupled DNA repair factor that directs RNA polymerase II ubiquitylation"

**This PDF file includes:**

Materials and Methods
Figs. S1 to S10
Tables S1 to S7

### Materials and Methods

**Cell lines.** All cell lines (listed in Table S1) were cultured at 37°C in an atmosphere of 5% CO2 in DMEM (Thermo Fisher Scientific) supplemented with penicillin/streptomycin (Sigma) and 10% fetal bovine serum (FBS; Bodinco BV or Thermo Fischer Scientific (Gibco)). U2OS Flp-In/T-REx cells (from here on out called U2OS (FRT)), which were generated using the Flp-InTM/T-RExTM system (Thermo Fisher Scientific), were a gift from Daniel Durocher (Panier et al., 2012).

**Construction of RPE1-TetOn-Cas9-PuroS-*TP53*-KO.** Human RPE1-hTERT cells were acquired from ATCC (CRL-4000™). Lentiviral particles of pLVX-Tet3G (Clontech) and pLVX-Tre3G-Cas9 plasmid (Clontech, Cas9 from vector Lenti-Cas9-2A-Blast (Addgene) were produced in HEK293T cells using the Lenti-X HT packaging system (Clontech). Transduced RPE1-hTERT cells were selected using 10 µg/mL puromycin and 400 µg/mL G418 to generate RPE1-TetOn-Cas9. To create knockouts, Cas9 expression was induced by 100 ng/mL doxycycline (Sigma-Aldrich) followed by transfection by RNAiMAX (Invitrogen) with 10 nM synthetic crRNA and 10 nM tracrRNA (Integrated DNA Technologies, IA). crRNAs were used the generate knockouts of *TP53* and the *Streptomyces* puromycin-N-acetyltransferase gene *PAC1* to re-introduce puromycin sensitivity. The population was selected for *TP53*-KO cells by selection on 10 µM Nutlin-3 (Selleck Chemicals, TX) for one week. Single cell clones were selected based on sensitivity to 1 µg/mL puromycin, and knockout of *TP53* was confirmed by Sanger sequencing and western blot. Cas9 activity of the RPE1-TetOn-Cas9-PuroS-*TP53*-KO cell line (RPE1-iCas9) was assayed by transducing cells with lentivirus produced using vector pXPR-011 (Addgene) (Doench et al., 2014) containing eGFP, sgGFP and Puromycin N-acetyltransferase. After transduction, transduced cells were selected by puromycin (3 µg/mL, Sigma) and eGFP-expression was assayed every 4 h for up to a week in the absence and presence of doxycycline (100ng/mL, Sigma) in an IncuCyte Zoom automated live-cell imaging platform (Sartorius, Göttingen, Germany).

**CRISPR/Cas9 screens.** For every screen, three populations of RPE1-iCas9 were transduced at an M.O.I. of ~0.2 with a 1:1000 dilution of TKOv3-in-pLCKO lentiviral library in medium containing 8µg/mL hexadimethrine bromide (Sigma-Aldrich). The library was a gift from Katherine Chan, Amy Tong and Jason Moffat (Donnelly Centre, University of Toronto, Toronto, ON). Twenty-four h after transduction, puromycin (Sigma-Aldrich) was added to 5 µg/mL to select for transduced cells. After selection was complete and all cells in non-transduced control populations had died, and dishes with transduced populations had reached 90% confluence, a t=0 sample was taken for each of the three populations. From the remaining cells of each population $30 \times 10^6$ (corresponding to a library representation of >400) were grown as a control population, and additionally $30 \times 10^6$ were grown in the presence of Illudin S for the drug screens, at a concentration of up to 25 nM for the screen in RPE1-iCas9, up to 2 nM for the screen in RPE1-

iCas9 *CSB*-KO, and up to 4 nM for the screen in RPE1-iCas9 *ELOF1*-KO. Doxycycline was added to the medium of all replicates from t=0 onwards to induce expression of Cas9, at a concentration of 200 ng/mL. After three doublings 30 × $10^6$ cells of each population were passed. After 12 doublings, all populations were harvested.

**Sequencing and analysis of CRISPR screens.** Genomic DNA was isolated from each population using the Blood and Cell Culture DNA Maxi Kit (Qiagen), dissolved in a 2mM Tris-Cl, 0.1 mM EDTA buffer (pH 9.0) and the concentrations were measured by NanoDrop spectrophotometer. Lentiviral inserts were amplified from the genomic DNA of each population in 64 parallel 50µl reactions, each reaction containing 3 µg gDNA in 24.5 µl buffer as template, 0.25 µl FW primer (TKO Outer Fw, 100µM), 0.25µl RV primer (TKO Outer Rv, 100µM) and 25 µl KAPA HiFi HotStart 2x ReadyMix (Roche). The PCR conditions were: 95°C for 3 min; 28 cycles at 98°C for 20 s, 66°C for 15 s and 72°C for 15 s; 72°C for 1 min; hold at 4°C. After the PCR, the 64 reactions of each sample were pooled, mixed, and 0.5µl was used as template for the second PCR reaction. In the second PCR reaction, reverse primers with different Illumina i7 index sequences were used for each sample to identify the sample after pooled sequencing as described by Hart *et al*. (Hart et al., 2015). Each reaction contained 0.5µl of the first PCR as template, 16.5µl H2O, 4µl FW primer (TKO Fw i5 index 1, 10µM), 4µl RV primer (One of twelve TKO Rv i7 index primers)) and 25µl KAPA HiFi HotStart ReadyMix (2x). PCR conditions were: 95°C for 3 min; 16 cycles at 98°C for 20 s, 66°C for 15 s and 72°C for 15 s; 72°C for 1 min; hold at 4°C. All oligonucleotides for screen sample preparation were obtained from Integrated DNA Technologies, the primers for the second PCR reaction were ordered as HPLC-purified oligos. The second PCR products of each pool were purified by QIAquick PCR Purification Kit (Qiagen), concentrations were determined by NanoDrop and samples were combined in an equimolar pool. RNAseq samples were added to 10% to create balanced reads for the first 21 nucleotides sequenced, as for all the screen samples these read the U6 promoter sequence upstream of the sgRNA sequence. Up to twelve samples were sequenced in a single HiSeq4000 lane in an SR50 run using standard reagents and conditions and reads were mapped to the TKOv3 library sequences, not allowing any mismatches. The screen data was analyzed using DrugZ version 1.1.0.2 (Colic et al., 2019), $Log_2$_Fold-Change was calculated by first normalizing all samples to an equal number of total reads (pseudocount +1), then calculating the median fold change per guide for the three replicate screen samples, and finally calculating the $Log_2$ of the median fold change of all guides targeting the gene. For network analysis, the physical interactions of differentially depleted genes were extracted from the gene interaction database, GeneMANIA. Functional pathway analysis was performed using Reactome. The interaction network was built and visualized in Cytoscape v.3.7. Node colors were adjusted based on FDR. Node border colors were customized based on enriched pathways.

**Generation of knock-out cells.** Cells were either transfected with Cas9-2A-GFP (pX458; Addgene #48138) containing a guide RNA from the TKOv3 library (Addgene #125517) or co-transfected with Cas9-2A-GFP (pX458; Addgene #48138) together with pLV-U6g-PPB encoding a guide RNA from the LUMC/Sigma-Aldrich sgRNA library (sgRNAs are listed in Table S2 and plasmids are listed in Table S4) using lipofectamine 2000 (Invitrogen). Cells were FACS sorted on BFP/GFP and plated at low density after which individual clones were isolated. Alternatively, cells were selected with puromycin (1 µg/mL) for 3 days and seeded at low density after which individual clones were isolated. Isolated knockout clones were verified by western blot analysis and/or Sanger sequencing.

**PCR analysis of knockout clones.** Genomic DNA was isolated by resuspending cell pellets in WCE buffer (50mM KCL, 10mM Tris pH 8.0, 25 mM MgCl2 0.1 mg/mL gelatin, 0.45% Tween-20, 0.45% NP-40) containing 0,1 mg/mL Proteinase K (EO0491;Thermo Fisher Scientific) and incubating for 1h at 56°C followed by a 10 min heat inactivation of Proteinase K by 96°C. Fragments of approximately 1kb, containing the sgRNA sequence, were amplified by PCR (sequencing primers are listed in Table S5) followed by Sanger sequencing using either the forward or the reverse primer.

**Plasmids.** The Neomycin resistance gene in pcDNA5/FRT/TO-Neo (Addgene #41000) was replaced with a Puromycin resistance gene. Fragments spanning GFP-N1 (Clontech) including the multiple cloning site were inserted into pcDNA5/FRT/TO-puro. $ELOF1^{WT}$ was amplified by PCR (Primers are listed in Table S6) and inserted into pcDNA5/FRT/TO-puro-GFP-N1. ELOF1 mutants were generated by site-directed mutagenesis PCR (Primers are listed in Table S6). All sequences were verified by Sanger sequencing.

**Generation of stable cell lines.** U2OS (FRT) ELOF1-KO clone 3-12 was selected and subsequently used to stably express inducible GFP-tagged proteins (Cell lines are listed in Table S1) by co-transfecting pCDNA5/FRT/TO-Puro plasmid encoding GFP-tagged fusion proteins (5 µg), together with pOG44 plasmid encoding the Flp recombinase (0.5 µg) using lipofectamine 2000 (Invitrogen). Cells expressing inducible GFP-tagged proteins were selected by incubation with 1 µg/mL puromycin and 4 µg/mL blasticidin S. Expression of these GFP-tagged proteins was induced by the addition of 2 µg/mL doxycycline for 24 h.

**Immunoprecipitation for Co-IP.** Cells were mock treated or irradiated with UV-C light (20 J/m$^2$) and harvested 1 h after UV. Chromatin-enriched fractions were prepared by incubating the cells for 20 min on ice in IP buffer (IP-130 for endogenous RNAPII IP and IP-150 for GFP-IP), followed by centrifugation, and removal of the supernatant. For endogenous RNA pol II IPs the chromatin-enriched cell pellets were lysed in IP-130 buffer (30 mM Tris pH 7.5, 130 mM NaCl, 2 mM MgCl2, 0.5% Triton X-100, protease inhibitor cocktail (Roche), 250 U/mL Benzonase® Nuclease (Novagen), and 2 µg RNAPII-S2 (ab5095, Abcam) for 2-3 h at 4 °C. For GFP IPs the chromatin-enriched cell pellets were lysed in IP-150 buffer (50 mM Tris pH 7.5, 150 mM NaCl, 0.5% NP-40, 2 mM $MgCl_2$, protease inhibitor cocktail (Roche), and 500 U/mL Benzonase® Nuclease (Novagen)) for 1 h at 4 °C. Protein complexes were pulled down by 1.5 h incubation with Protein

A agarose beads (Millipore) or GFP-Trap® A beads (Chromotek). For subsequent analysis by western blotting, the beads were washed 6 times with IP-130 buffer for endogenous RNAPII IPs and EBC-2 buffer (50 mM Tris pH 7.5, 150 mM NaCl, 1 mM EDTA, 0.5% NP-40, and protease inhibitor cocktail (Roche)) for GFP-IPs. The samples were prepared by boiling in Laemmli-SDS sample buffer. Bound proteins were separated by sodium dodecyl sulfate polyacrylamide gel electrophoresis (SDS-PAGE) and immunoblotted with the indicated antibodies.

**Protein degradation.** RPE1-iCas9 cells were mock treated or irradiated with UV-C light (9 J/m$^2$) and incubated in conditioned media for different periods of time (7, 24, 48h). Cells were harvested by scraping in Laemmli-SDS sample buffer and proteins were separated by SDS-PAGE and immunoblotted with the indicated antibodies.

**Western blotting.** Proteins were separated on 4-12% Criterion XT Bis-Tris gels (Bio-Rad, #3450124) in NuPAGE MOPS running buffer (NP0001-02 Thermo Fisher Scientific), and blotted onto PVDF membranes (IPFL00010, EMD Millipore). Membranes were blocked with blocking buffer (Rockland, MB-070-003) for 1 h at RT. Membranes were then probed with antibodies (Antibodies are listed in Table S3) as indicated. Proteins stained with HRP-conjugated secondary antibodies were detected using Western Lightning Plus-ECL (PerkinElmer, NEL103001EA).

**Clonogenic survival assays.** Knockout and rescue cell lines were trypsinized, seeded at low density and mock-treated or exposed to a dilution series of Illudin S (Santa Cruz; sc-391575) for 72 h (50 and 100 pg/mL or 50 and 200 pg/mL). On day 10, the cells were washed with 0.9% NaCl and stained with methylene blue. Colonies of more than 20 cells were scored.

**Propidium-iodide (PI) proliferation assay.** 350 cells per well were seeded in a 384-well plate, and exposed to Illudin S. After 72 h drug exposure, cells were fixed for 1 h in 4% paraformaldehyde, permeabilized and stained using 0.1% Tween-20 and 0.1mg/mL of propidium iodide in PBS, and imaged by IncuCyte Zoom using a 4x objective.

**Dose-response curves.** Cells were seeded in 96-well plates (1000 cells per well for RPE1-iCas9) in 150 µl of medium. 24 h after seeding, drugs were added manually dissolved in 50 µl of medium (Cisplatin) or printed directly into the plate by Tecan D300e digital dispenser (Illudin S, Irofulven). Phenylarsine oxide (PAO; Sigma-Aldrich) was used at 10 µM as a control for zero cell viability. After 72 h of drug exposure, cell viability was assayed by adding 30 µL of CellTiter-Blue (Promega) in medium to each well. Plates were incubated for 4 h at 37°C, and plate fluorescence (560Ex/590Em) was measured on a BioTek plate reader. Relative viability was calculated for each well from the fluorescence measurements as: (value-(value(PAO))/((value(untreated)-(value(PAO)). A four-parameter dose–response curve was fit to the viability values by Graphpad Prism 7.

**RNA recovery synthesis (RRS).** Cells were seeded on 12-mm coverslips in DMEM supplemented with 10% FBS. Medium was changed to DMEM supplemented with 1% FBS for at least 24 h prior to the nascent transcript measurement to reduce the excess of available uridine in the culture medium. For the RRS following DRB treatment, cells were supplemented with 100 µM DRB (Sigma, D1916) for 2 h and either during the DRB treatment or following DRB washout the cells were pulse-labelled with 400 µM 5-ethynyl-uridine (EU; Jena Bioscience) for 1 h. The cells were washed with in phosphate-buffered saline (PBS), fixed with 3.7% formaldehyde in PBS for 15 min, and stored in PBS at 4°C. For the RRS following UV-irradiation, cells were UV-irradiated (9 $J/m^2$ or 12 $J/m^2$), allowed to recover for the indicated time periods, and pulse-labelled with 400 µM 5-ethynyl-uridine (EU; Jena Bioscience) for 1 h. After medium-chase with DMEM without supplements for 15 min, cells were fixed with 3.7% formaldehyde in PBS for 15 min and stored in PBS at 4°C. Cells were permeabilized with 0.5% Triton X-100 in PBS for 10 min at RT and blocked in 1.5% bovine serum albumin (BSA, Thermo Fisher) in PBS. Nascent RNA was visualized by click-it chemistry, labeling the cells for 1 h with a mix of 60 µM Atto azide-Alexa594 (Atto Tec), 4 mM copper sulfate (Sigma), 10 mM ascorbic acid (Sigma) and 0.1 µg/mL DAPI in a 50 mM Tris-buffer. Cells were washed extensively with PBS and mounted in Polymount (Brunschwig).

**Unscheduled DNA synthesis (UDS).** Cells were seeded on 18-mm coverslips in DMEM with 1% FBS. After 24 h, cells were locally irradiated through a 5 µm filter with 30 $J/m^2$ UV-C. Cells were subsequently pulse-labelled with 20 µM 5-ethynyl deoxy-uridine (EdU; VWR) and 1 µM FuDR (Sigma Aldrich) for either 1 h or 4 h. After medium-chase with DMEM without supplements for 30 min, cells were fixed with 3.7% formaldehyde in PBS for 15 min and stored in PBS at 4°C. Cells were permeabilized with 0.5% Triton X-100 in PBS for 20 min at RT and blocked in 3% BSA (Thermo Fisher) in PBS. The incorporated EdU was coupled to Atto azide Alexa Fluor 647 using Click-iT chemistry according to the manufacturer's instructions (Invitrogen). After coupling, the cells were post-fixed with 2 % formaldehyde for 10 min and subsequently blocked with 100 mM Glycine. DNA was denatured with 0.5 % NaOH for 5 min, followed by blocking with 10 % BSA (Thermo Fisher) for 15 min. Next, the cells were incubated with an antibody against CPDs (Antibodies are listed in Table S3) for 2 h, followed by secondary antibodies 1 h, and DAPI for 5 min. Cells were mounted in Polymount (Brunschwig).

**Microscopic analysis of fixed cells.** Images of fixed samples were acquired on a Zeiss AxioImager M2 or D2 widefield fluorescence microscope equipped with 63x PLAN APO (1.4 NA) oil-immersion objectives (Zeiss) and an HXP 120 metal-halide lamp used for excitation. Fluorescent probes were detected using the following filters: DAPI (excitation filter: 350/50 nm, dichroic mirror: 400 nm, emission filter: 460/50 nm), Alexa 555 (excitation filter: 545/25 nm, dichroic mirror: 565 nm, emission filter: 605/70 nm), Alexa 647 (excitation filter: 640/30 nm, dichroic mirror: 660 nm, emission filter: 690/50 nm). Images were recorded using ZEN 2012 software and analyzed in Image J.

**ChIP-sequencing.** Cells were mock treated or UV-irradiated with 9 J/m$^2$ and incubated in conditioned media for different periods of time (1, 4, 8, 16 h). Cells were crosslinked with 0.5 mg/mL disuccinimidyl glutarate (DSG; Thermo Fisher) in PBS for 45 min at room temperature. Cells were washed with PBS and crosslinked with 1% PFA for 20 min at room temperature. Fixation was stopped by adding 1.25 M Glycin in PBS to a final concentration of 0.1 M for 3 minutes at room temperature. Cells were washed with cold PBS and lysed and collected in a buffer containing 0.25% Triton X-100, 10 mM EDTA (pH 8.0), 0.5 mM EGTA (pH 8.0) and 20 mM Hepes (pH 7.6). Chromatin was pelleted in 5 min at 400 g and incubated in a buffer containing 150 mM NaCl, 1 mM EDTA (pH 8.0), 0.5 mM EGTA (pH 8.0) and 50 mM Hepes (pH 7.6) for 10 minutes at 4°C. Chromatin was again pelleted for 5 min at 400 g and resuspended in ChIP-buffer (0.15 % SDS, 1 % Triton X-100, 150 mM NaCl, 1 mM EDTA (pH 8.0), 0.5 mM EGTA (pH 8.0) and 20 mM Hepes (pH 7.6)) to a final concentration of 15x10$^6$ cells/mL. Chromatin was sonicated to approximately 1 nucleosome using a Bioruptor waterbath sonicator (Diagenode). Chromatin of ~5x10$^6$ cells was incubated with 3 µg antibody (Antibodies are listed in Table S3) overnight at 4°C, followed by a 1.5 h protein-chromatin pull-down with a 1:1 mix of protein A and protein G Dynabeads (Thermo Fisher; 10001D and 10003D). ChIP samples were washed extensively, followed by decrosslinking for 4 h at 65°C in the presence of proteinase K. DNA was purified using a Qiagen MinElute kit. For ser2-RNAPII ChIP-seq sample libraries were prepared from 1 ng ChIPed DNA using NEBNext Ultra II DNA Library Prep Kit for Illumina (NEB, E7645S) and index primer sets (NEB, E7335S). Strand-biased library amplification was performed using high-fidelity KAPA HiFi Enzyme (Roche, KK2102). The prepared libraries were sequenced on an Illumina HiSeq 2500 system (Illumina), resulting in 150 bp paired-end reads. For the remaining ChIP-seq samples, libraries were prepared using KAPA HyperPrep kit (Roche) and A-T mediated ligation of full Y-shaped IDT adapters. Samples were sequenced in a 150bp paired-end run on an Illumina HiSeq X system (Macrogen).

**BrU-sequencing.** For BrU-sequencing after DRB treatment, cells were pre-treated with DRB (100 µM) for 1 h. Then cells were washed with PBS and incubated with 2 mM bromouridine (Bru) at 37°C for 15 or 30 min, as indicated per experiment. For BrU sequencing after UV-irradiation, cells were UV-irradiated with 9J/m$^2$. Cells were then incubated in conditioned media for different periods of time (3, 8, 24h) before being incubated with 2 mM bromouridine (Bru) at 37°C for 30 min. All BrU-sequencing samples were subsequently lysed in TRIzol reagent (Invitrogen) and Bru-containing RNA was isolated as previously described (Andrade-Lima et al., 2015). cDNA libraries were made from the Bru-labeled RNA using the Illumina TruSeq library kit and paired-end 151 bp sequenced using the Illumina NovaSeq platform at the University of Michigan Advanced Genomics Core. Only single-end sequencing data was used for downstream analyses.

**ATAC-seq.** For each experimental condition 50.000 cells were harvested and washed with cold PBS containing protease inhibitors (Roche). The cells were resuspended in 50 µl Transposase mixture, containing 1x Tagment DNA buffer Illumina Kit, #20034197), 0.5 µl Tagment DNA enzyme (Illumina Kit, #20034197) and 200 ng/µl Digitonin (Promega, #G9441), and incubated at 37 °C for 30 min. Fragmented DNA was purified using a Qiagen MinElute kit (#28204). Libraries were prepared by PCR using NEBNext High-Fidelity 2x PCR Master Mix (New England Labs, #M0541) and Nextera XT index Kit v2 (Illumina). Samples were sequenced in a 150bp paired-end run on an Illumina HiSeq X system (Macrogen).

**RNA-seq and data analysis.** Cells were UV-irradiated (254 nm, 9 J/m$^2$) or mock-treated and harvested 24 h later. Total RNA was isolated using the RNeasy mini kit (Qiagen) according to the manufacturer's protocol. Up to $5 \times 10^6$ cells per sample were lysed in RLT buffer. For quantification purposes, 24 ng of the eight ArrayControl™ RNA Spikes (Thermo Fisher Scientific) were added to each sample at this stage (3 ng/spike). Samples were purified using the KAPA mRNA HyperPrep kit (Roche) and prepared for sequencing using the TruSeq RNA Library Prep Kit v2 (Illumina) according to the manufacturer's instructions, and sequenced in an SR50 run in a single lane on an Illumina HiSeq4000. For RNA-seq data analysis, obtained sequencing reads were cleaned by 5'-end quality trimming and Illumina-adapter clipping by Trimmomatic (Bolger et al., 2014). Pre-alignment quality control of the cleaned sequencing reads was done with FastQC.The alignment to reference genome hg19 of trimmed sequencing reads was done with Hisat2, guided by gene annotation in the refGene UCSC table (Kim et al., 2015). The generated SAM files were converted to the binary counterpart BAM, followed by BAM sorting and indexing with SAMtools (Li et al., 2009). Counting reads to the UCSC's genomic features hg19, performed by Subread and featureCounts (*8*). Differential expression analysis was performed using edgeR (Robinson et al., 2010). Only genes with at least 2 counts per million in at least 33% of samples were included in the analysis. Data were normalized for sample specific effects by the trimmed mean of M-values. This was followed by estimating the dispersion and determining the differentially expressed genes, using general linear model (GLM). FDR-adjusted p-values < 0.05 were considered significant.

**ChIP-seq, BrU-seq and ATAC-seq data analyses.** For ser2-RNAPII ChIP-seq, low-quality sequence reads and adapters were filtered out by Trimmomatic (v3.36) (Bolger et al., 2014). The trimmed reads were aligned to the human reference genome (GRC h37/hg19) with the Burrows-Wheeler Aligner (BWA-v0.7.12-r1039) (Li, 2013). Biobambam2 (v2.0.72) (Tischler and Leonard, 2014) was used to remove duplicate reads from the aligned reads. Sequence reads were locally realigned and base-quality scores were recalibrated with the IndelRealigner and BaseRecalibrator programs in Genome Analysis Toolkit (GATK-v3.5) (McKenna et al., 2010).

For pan-RNAPII ChIP-seq, BrU-seq and ATAC-seq, reads were aligned to the Human Genome 38 (Hg38) using bwa-mem tools (BWA (Version 0.7.16a) or STAR (Version 2.5.3a)) (Dobin et al., 2013; Li, 2013). Only high-quality reads (> q30) were included in the analyses and duplicates were removed using Samtools (Version 1.6) with fixmate -m and markdup -r settings (Table S7). Bam files were converted into stranded TagDirectories and UCSC genome tracks using HOMER tools (Version 4.8.2) (Heinz et al., 2010). A list of 49,948 genes was obtained from the

UCSC genome database (https://genome.ucsc.edu/cgi-bin/hgTables) selecting the knownCanonical table containing the canonical transcription start sites per gene. To prevent contamination of binding profiles, genes should be non-overlapping with at least 2 kb between genes. Subsequent selection of sets of genes is described per analysis.

For ChIP-seq, binding profiles within selected areas of individual genes (e.g. around TSS or TTS), were defined using the AnnotatePeaks.pl tool of HOMER using the default normalization to 10 million reads. Metagene profiles were defined using the makeMetaGeneProfile.pl tool of HOMER, using default settings. Read densities in input samples were subtracted from individual ChIP-seq datasets to background-correct our data in which negative values were converted to 0, to prevent the use of impossible negative read densities in further calculations. Individual datasets were subsequently processed into heatmaps or binding profiles using R (Version 3.5.3) (Team, 2019). ChIP-seq binding and metagene profiles were averaged per set of genes and profiles were normalized to area under the curve to allow proper comparison of the profiles without effects of overall differences in read density.

ATAC-seq and BrU-seq heatmaps were generated as described for ChIP-seq analyses, except for the input subtraction. BrU-seq aggregated profiles of Figure 3D and Figure S4B, were defined using the AnnotatePeaks.pl tool of HOMER using the default normalization to 10million reads, as for ChIP-sequencing. Genes were selected on size and strongest binding of RNAPII at the TSS in undamaged condition, as for ChIP-sequencing. Averaged binding profiles were subsequently normalized to nascent transcript levels, as quantified by 5-EU labelling, relative to the control in their specific cell type. For example, WT cells 3 h after UV irradiation showed 35% RNA relative to WT control cells, so we multiplied the expression of the bins by 0.35. For Figure 2J and Figure S3D, genes were selected based on their lengths for the different analyses (either 25-50 kb, 50-100 kb, or greater than 100 kb), TSSs were at least 10 kb apart and expression was at least 0.05 RPKM. Base coverages were used to compute read counts TSS or TTS which were then normalized to feature length and number of uniquely-mapped reads (RPKM method).

**Strand-bias analyses and recovery index.** Pull down of RNAPII from damaged genes would result in a relative overrepresentation of damaged transcribed strands in the ChIP-seq samples. The inability of high-fidelity DNA polymerases to amplify damaged DNA strands, therefore, results in unequal PCR amplification of the transcribed and non-transcribed strand during ChIP-seq library sample prep. ChIP-seq reads were quantified in transcribed and non-transcribed strands of genes of 3-100 kb with minimal gap of 2 kb between genes using the AnnotatePeaks.pl tool of HOMER with the default normalization to 10 million reads. We focused on the top 3000 genes that showed strongest binding of RNAPII at the TSS in WT undamaged condition to select for actively transcribed genes. A strand specificity index (SSI) was subsequently calculated per gene with the following equation (as described before (Nakazawa et al., 2020))

$$\text{Strand specificity index} = \frac{\text{reads in "transcribed" strand} - \text{reads in "non} - \text{transcribed" strand}}{\text{reads in "transcribed" strand} + \text{reads in "non} - \text{transcribed" strand}}$$

Depending on whether genes are present on the + or – strand of DNA, a strand bias results in either positive or negative SSI values, varying around 0 (representing no strand bias).

To quantify the amount of damage and correlated level of recovery, we defined a recovery index (RI). For this, we made frequency distributions of per-gene SSIs. While in non-biased ChIP-sequencing samples SSIs frequency distributions would follow a single Gaussian distribution, SSI frequency distribution of biased ChIP-seq samples are expected to follow a mixed Gaussian distribution composed of 3 normal distributions; one distribution representing genes without strand bias (with $\mu_0$=0), and two Gaussian distributions representing strand-biased genes on either the + or – strands of DNA, with $\mu_1$ and $\mu_2$ symmetrically positioned around $\mu_0$ ($-\mu_1 = \mu_2$). The mean distance that the strand-biased Gaussian distributions deviate from 0 (which is represented by $-\mu_1$ or $\mu_2$) was defined as the recovery index (RI). The three Gaussian distributions were fitted using normalmixEM of the mixtools package in R (Benaglia T, 2009).

**Data and materials availability:** Both raw and processed ChIP-seq, BrU-seq, ATAC-seq, and RNA-seq data shown in main Figure 2, 3, 5, and Fig S4 – S9 are deposited in the Gene Expression Omnibus (GEO) under GSE149760 (password: obmhawiqvnenxal). Additional data and custom code will be made available upon reasonable request.

### Table S1. Cell lines

| Cell line | Source |
|---|---|
| U2OS(FRT) | (Panier et al., 2012) |
| U2OS(FRT) CSB-KO (1-12) | (van der Weegen et al., 2020) |
| U2OS(FRT) ELOF1-KO (3-12) | This study |
| U2OS(FRT) ELOF1-KO (3-12) + ELOF1-GFP-N1 | This-study |
| U2OS(FRT) ELOF1-KO (3-12) + ELOF1ΔN-GFP | This-study |
| U2OS(FRT) ELOF1-KO (3-12) + ELOF1S72/D73K-GFP | This-study |
| RPE1-iCas9 | This study |
| RPE1-iCas9 CSB-KO (1-15) | This-study |
| RPE1-iCas9 ELOF1-KO (2-12) | This-study |
| RPE1-iCas9 ELOF1-KO (2-16) | This-study |
| RPE1-iCas9 UVSSA-KO (3-9) | This-study |
| RPE1-iCas9 PTGR1-KO (11) | This-study |
| RPE1-iCas9 XPC-KO | This-study |
| RPE1-iCas9 CSB-KO (15)/ELOF1-KO (2-20) | This-study |
| RPE1-iCas9 CSB-KO (15)/CSA-KO (3-21) | This-study |

### Table S2. sgRNAs

| sgRNA | Sequence | Gene_ID | Exon_ID | Exon |
|---|---|---|---|---|
| CSA_3 | GGAGAGCAGAGTCAACACGG | ENSG00000049167 | ENSE00001762056 | 1 |
| CSB_1 | CTCATCGGATCATTCTGTCT | ENSG00000225830 | ENSE00002514316 | 10 |
| UVSSA_3 | GCCGGCTGTGTGCTCGTGGA | ENSG00000163945 | ENSE00001713804 | 6 |
| ELOF1_2 | CGAGAAATCCTGTGATGTGA | ENSG00000130165 | ENSE00003469475 | 2 |
| ELOF1_3 | TGATTGGATAGACGCCTGCG | ENSG00000130165 | ENSE00001061357 | 4 |
| PTGR1_1 | CGGTGAGGAACAAAGCTTCA | ENSG00000106853 | ENSE00003807576 | 3 |
| PTGR1_2 | GTAGGCCAAAGTAGGCAGTC | ENSG00000106853 | ENSE00003810161 | 6 |
| TP53 | CCATTGTTCAATATCGTCCG | ENSG00000141510 | ENSE00003625790 | 4 |
| PAC1 | ACGCGCGTCGGGCTCGACAT | | | |

#### Table S3. Antibodies

| Antibody | Host | Company (reference) | Use | Identifier |
|---|---|---|---|---|
| Cas9 | Mouse | Cell signalling, #14697 (7A9-3A3) | WB: 1:2000 | aML#031 |
| CSA/ERCC8 | Mouse | Santa Cruz, #sc-376981 (D2) | WB: 1:500 | aML#025 |
| CSA/ERCC8 | Rabbit | Abcam, #137033 (EPR9237) | WB: 1:500 | aML#028 |
| CSB/ERCC6 | Goat | Santa Cruz, #sc-10459 (E-18) | WB: 1:1000 | aML#039 |
| CSB/ERCC6 | Rabbit | Santa Cruz, #sc-25370 (H-300) | WB: 1:300 | aML#003 |
| CSB/ERCC6 | Mouse | Bio Matrix Research, #BMR00638 (553C5a) | WB: 1:1000 | |
| GFP | Mouse | Roche, #11814460001 (7.1 and 13.1) | WB: 1:1000 | aML#011 |
| GFP | Rabbit | Abcam, #ab290 | WB: 1:1000 | aML#044 |
| GFP | Mouse | Santa Cruz, #sc-9996 (B2) | WB: 1:1000 | |
| p62/GTF2H1 | Mouse | Santa Cruz, #sc-48431 (G10) | WB: 1:500 | aML#099 |
| p89/XPB/ERCC3 | Mouse | Millipore, #MABE1123 | WB: 1:2000 | aML#101 |
| p89/XPB/ERCC3 | Mouse | Santa Cruz, #sc-271500 (G10) | WB: 1:1000 | |
| UVSSA | Mouse | Abnova, #H00057654-B01P | WB: 1:500 | |
| RNAPII-S2 | Rabbit | Abcam, #ab5095 | WB: 1:1000 | aML#024 |
| RNAPII-S2 | Rat | Milipore, #04-1571 (3E10) | ChIP: 3µg, WB: 1:1500 | aML#120 |
| RNAPII-Total | Rabbit | Bethyl, #A304-405A | ChIP: 3µg | aML#088 |
| ATF3 | Rabbit | Abcam, #ab207434 (EPR19488) | WB: 1:1000 | aML#064 |
| γH2aX | **Mouse** | Millipore, #05-636 (JBW301) | WB: 1:2500 | |
| Flag | Mouse | Sigma, #F1804 (M2) | WB: 1:5000 | |
| Tubulin | Mouse | Sigma, #T6199 (DM1A) | WB: 1:1000 | aML#008 |
| Mouse IgG (H+L) CF770 | Goat | Biotium, VWR #20077 | WB: 1:10000 | aML#009 |
| rabbit IgG (H+L) CF680 | Goat | Biotium, VWR #20067 | WB: 1:10000 | aML#010 |
| Goat IgG (H+L) CF680 | Donkey | Thermo fisher Scientific, #A21084 | WB: 1:10000 | aML#037 |
| Mouse IgG (HRP) | Goat | Abcam, #ab6789 | WB: 1:5000 | aML#132 |
| Rabbit IgG (HRP) | Goat | Abcam, #ab6721 | WB: 1:5000 | aML#105 |
| Rat IgG (HRP) | Goat | Cell signaling, #7077 | WB: 1:1000 | |
| Mouse IgG (HRP) | Horse | Cell signaling, #7076 | WB: 1:1000 | |
| Mouse Alexa 555 | Goat | Therme fisher Scientific, A-21424 | IF: 1:1000 | aML#015 |
| CPD | Mouse | Cosmo Bio, CAC-NM-DND-001 | IF: 1:1500 | aML#020 |

### Table S4. Plasmids

| Plasmid | Origin |
|---|---|
| pcDNA5/FRT/TO-Neo | Addgene #41000 |
| pcDNA5/FRT/TO-Puro | This study |
| pcDNA5/FRT/TO-Puro-GFP-N1 | This study |
| pLV-U6g-PPB | LUMC/Sigma-Aldrich sgRNA library |
| pOG44 | Thermo Fisher |
| pX458 | Addgene #48138 |
| pXPR_011 | Addgene #59702 |
| pX458-(Cas9-2A-GFP)-sgELOF1-2 | This study |
| pX458-(Cas9-2A-GFP)-sgELOF1-3 | This study |
| pX458-(Cas9-2A-GFP)-sgUVSSA_3 | This study |
| pX458-(Cas9-2A-GFP)-sgERCC8_3 | This study |
| pcDNA5/FRT/TO-puro-ELOF1-GFP-N1 | This study |
| pcDNA5/FRT/TO-puro-ELOF1ΔN-GFP-N1 | This study |
| pcDNA5/FRT/TO-puro-ELOF1-S72K-D73K-GFP-N1 | This study |

### Table S5. Sequencing primers

| Gene | Sequence | Identifier |
|---|---|---|
| CSB | 5-GTAGGGGCCAGTTGTTAGAATGTAA-3 | oML#078_sgML#003_CSB1_fw |
| | 5-CTCACATTCTGAATGACTTGGCTA-3 | oML#079_sgML#003_CSB1_rev |
| CSA | 5-ACTGACCTCGCAATCACTGAC-3 | oML#444_pML#219_CSA3FWa |
| | 5-CAAGTACACAAGGCTCTTCCTC-3 | oML#445_pML#219_CSA3RVa |
| | 5-TTGGTCCGTGCCCACGTG-3 | oML#446_pML#219_CSA3FWb |
| | 5-GTCCTCTGCCTTTAATAGGCTG-3 | oML#447_pML#219_CSA3RVb |
| UVSSA | 5-ACCCAGAGGTACACAGAGATTG-3 | oML#090_sgML#019_UVSSA1_Fw |
| | 5-GCTCTTAGAAGTGTCCCTGTG-3 | oML#091_sgML#019_UVSSA1_Rv |
| | 5-ATCAGGAGGCTGAGGCGGCTG-3 | oML#076_sgML#020_UVSSA2_fw |
| | 5-AGGAGCCTACCCGGGAGCCGGG-3 | oML#077_sgML#020_UVSSA2_rev |
| ELOF1 | 5-ATGTTGCCCAGGCTGGTATC-3 | oML#320_sgELOF1-2_Seq_FW |
| | 5-TCCTCTGTGTCGCTACTGATTG-3 | oML#321_sgELOF1-2_Seq_RV |
| | 5-GATCACAGGTGTGAGCCAC-3 | oML#328_sgELOF1-2_seq_FW2 |
| | 5-CACTTAGGTCAAGGGCGATC-3 | oML#329_sgELOF1-2_seq_RV2 |
| | 5-AAGAAGATGACAGGCACCCTC-3 | oML#322_sgELOF1-3_Seq_FW |
| | 5-CGGGAAGTCCAGTTGAGATG-3 | oML#323_sgELOF1-3_Seq_RV |
| | 5-TGAAGGCGTCATCACCCAC-3 | oML#330_sgELOF1-3_seq_FW2 |
| | 5-GCTTTCGGAGCCAAGTGAG-3 | oML#331_sgELOF1-3_seq_RV2 |
| PTGR1 | 5-CCTCTCATGCCTAATATCACA-3 | oML#363_PTGR1_Fw2 |
| | 5-AGTACAACAGCCTTGGCAGT-3 | oML#364_PTGR1_Rv2 |
| | 5-GAACTTATATGGGAATGAGGCACA-3 | oML#369_PTGR1_Fw2-nested |
| | 5-GGGTGATGACTCTTAGCTGGCT-3 | oML#370_PTGR1_Rv2-nested |
| CRISRP_1 | AGGGCCTATTTCCCATGATTCCTT | TKO Outer Fw |
| | TCAAAAAAGCACCGACTCGG | TKO Outer Rv |
| CRISPR_2 | AATGATACGGCGACCACCGAGATCTACACTATAGCC TACACTCTTTCCCTACCGACGCTCTTCCGATCTTGTG GAAGGACGAGGA*C*C*G | TKO Fw i5 index 1<br><br>* phosphorothioate bonds |
| | CAAGCAGAAGACGGCATACGAGAT-(reverse-complement of i7 index)-GTGAC TGGAGTTCAGACGTGTGCTCTTCCGATCTATTTTAAC TTGCTATTTCTAGCTCTAA*A*A*C | TKO Rv i7 index primers<br><br>* phosphorothioate bonds |

### Table S6. Primers for cloning

| Gene | Sequence | Identifier |
|---|---|---|
| ELOF1-WT | 5-ACAATTGCTAGCGCCACCATGGGGCGCAGAAAGTC-3 | oML#308 |
| | 5-AGCTGTGTAAACCGGTAGCTGATTGGCCGCCTCG-3 | oML#309 |
| ELOF1ΔN | 5-GGTGTAACGGCTAGCGCCACCATGCCGCCTCCCAAGAAGAAGATG-3 | oML#345 |
| | 5-CCTGTCATCTTCTTCTTGGGAGGCGGCATGGTGGCGCTAGCCGTTA-3 | oML#346 |
| ELOF1S72/D73K | 5-GAACCCGTGGATGTGTACAAGAAGTGGATAGACGCCTGCGAG-3 | oML#353 |
| | 5-CTCGCAGGCGTCTATCCACTTCTTGTACACATCCACGGGTTC-3 | oML#354 |

### Table S7. Sequence depth of individual ChIP-seq, BrU-seq, RNA-seq replicates

| Technique | Cells | Condition | Sample names individual replicates | Unique, deduplicated reads |
|---|---|---|---|---|
| ATAC-seq | RPE1-iCas9 WT | WT mock-treated (n=2) | ATACseq_WTMock_Rep1_TagDir | 35,639,384 |
| | | | ATACseq_WTMock_Rep2_TagDir | 29,958,690 |
| | RPE1-iCas9 ELOF1-KO | ELOF1-KO mock-treated (n=2) | ATACseq_ELOFKOMock_Rep1_TagDir | 28,438,534 |
| | | | ATACseq_ELOFKOMock_Rep2_TagDir | 33,176,750 |
| BrU(DRB)-seq | RPE1-iCas9 WT | WT 15 minutes DRB release (n=1) | BrUseqRep2_WT15m_SSPE_TagDir | 38,261,822 |
| | | WT 30 minutes DRB release (n=1) | BrUseqRep2_WT30m_SSPE_TagDir | 36,625,983 |
| | RPE1-iCas9 ELOF1-KO | ELOF1-KO 15 minutes DRB release (n=1) | BrUseqRep2_ELOFKO15m_SSPE_TagDir | 29,354,235 |
| | | ELOF1-KO 30 minutes DRB release (n=1) | BrUseqRep2_ELOFKO30m_SSPE_TagDir | 36,297,661 |
| BrU-seq | RPE1-iCas9 WT | WT mock-treated (n=1) | BrUseqRep2_WT0h_SSPE_TagDir | 46,498,167 |
| | | WT 3 hours after 9J/m2 (n=1) | BrUseqRep2_WT3h_SSPE_TagDir | 30,486,325 |
| | | WT 8 hours after 9J/m2 (n=1) | BrUseqRep2_WT8h_SSPE_TagDir | 30,832,534 |
| | | WT 24 hours after 9J/m2 (n=1) | BrUseqRep2_WT24h_SSPE_TagDir | 46,960,735 |
| | RPE1-iCas9 ELOF1-KO | ELOF1-KO mock-treated (n=1) | BrUseqRep2_ELOFKO0h_SSPE_TagDir | 43,875,250 |
| | | ELOF1-KO 3 hours after 9J/m2 (n=1) | BrUseqRep2_ELOFKO3h_SSPE_TagDir | 34,909,758 |
| | | ELOF1-KO 8 hours after 9J/m2 (n=1) | BrUseqRep2_ELOFKO8h_SSPE_TagDir | 35,829,345 |
| | | ELOF1-KO 24 hours after 9J/m2 (n=1) | BrUseqRep2_ELOFKO24h_SSPE_TagDir | 35,592,827 |
| RNAPII-S2 ChIP-seq | RPE1-iCas9 WT | WT mock-treated (n=2) | DataTOgi_WTrep1_mock_SSPE_TagDir | 37,516,272 |
| | | | DataTOgi_WTrep2_mock_SSPE_TagDir | 38,571,922 |
| | | WT 1 hour after 9J/m2 (n=2) | DataTOgi_WTrep1_1h_SSPE_TagDir | 34,409,419 |
| | | | DataTOgi_WTrep2_1h_SSPE_TagDir | 38,789,028 |
| | | WT 4 hours after 9J/m2 (n=2) | DataTOgi_WTrep1_4h_SSPE_TagDir | 28,093,367 |
| | | | DataTOgi_WTrep2_4h_SSPE_TagDir | 32,867,919 |
| | | WT 8 hours after 9J/m2 (n=2) | DataTOgi_WTrep1_8h_SSPE_TagDir | 33,947,077 |
| | | | DataTOgi_WTrep2_8h_SSPE_TagDir | 17,812,398 |
| | | WT 16 hours after 9J/m2 (n=2) | DataTOgi_WTrep1_16h_SSPE_TagDir | 35,102,378 |
| | | | DataTOgi_WTrep2_16h_SSPE_TagDir | 45,062,679 |
| | RPE1-iCas9 CSB-KO | CSB-KO mock-treated (n=2) | DataTOgi_CSBKOrep1_mock_SSPE_TagDir | 37,986,546 |
| | | | DataTOgi_CSBKOrep2_mock_SSPE_TagDir | 40,640,212 |
| | | CSB-KO 1 hour after 9J/m2 (n=2) | DataTOgi_CSBKOrep1_1h_SSPE_TagDir | 38,447,432 |
| | | | DataTOgi_CSBKOrep2_1h_SSPE_TagDir | 36,096,454 |
| | | CSB-KO 4 hours after 9J/m2 (n=2) | DataTOgi_CSBKOrep1_4h_SSPE_TagDir | 33,023,511 |
| | | | DataTOgi_CSBKOrep2_4h_SSPE_TagDir | 34,115,863 |
| | | CSB-KO 8 hours after 9J/m2 (n=2) | DataTOgi_CSBKOrep1_8h_SSPE_TagDir | 38,924,552 |
| | | | DataTOgi_CSBKOrep2_8h_SSPE_TagDir | 39,151,481 |
| | | CSB-KO 16 hours after 9J/m2 (n=2) | DataTOgi_CSBKOrep1_16h_SSPE_TagDir | 34,581,457 |
| | | | DataTOgi_CSBKOrep2_16h_SSPE_TagDir | 43,678,724 |
| | RPE1-iCas9 ELOF1-KO | ELOF1-KO mock-treated (n=2) | DataTOgi_ELOFKOrep1_mock_SSPE_TagDir | 40,784,609 |
| | | | DataTOgi_ELOFKOrep2_mock_SSPE_TagDir | 43,805,221 |
| | | ELOF1-KO 1 hour after 9J/m2 (n=2) | DataTOgi_ELOFKOrep1_1h_SSPE_TagDir | 35,849,282 |
| | | | DataTOgi_ELOFKOrep2_1h_SSPE_TagDir | 46,626,951 |
| | | ELOF1-KO 4 hours after 9J/m2 (n=2) | DataTOgi_ELOFKOrep1_4h_SSPE_TagDir | 34,899,234 |
| | | | DataTOgi_ELOFKOrep2_4h_SSPE_TagDir | 43,369,451 |
| | | ELOF1-KO 8 hours after 9J/m2 (n=2) | DataTOgi_ELOFKOrep1_8h_SSPE_TagDir | 51,291,185 |
| | | | DataTOgi_ELOFKOrep2_8h_SSPE_TagDir | 9,676,403 |

| Technique | Cells | Condition | Sample names individual replicates | Unique, deduplicated reads |
|---|---|---|---|---|
| | | ELOF1-KO 16 hours after 9J/m2 (n=2) | DataTOgi_ELOFKOrep1_16h_SSPE_TagDir | 39,626,707 |
| | | | DataTOgi_ELOFKOrep2_16h_SSPE_TagDir | 42,524,194 |
| pan-RNAPII ChIP-seq | RPE1-iCas9 WT | WT input (n=2) | WT_input_rep1_SPPE_TagDir | 59,447,372 |
| | | | WT_input_rep2_SSPE_TagDir | 75,872,035 |
| | | WT mock-treated (n=2) | WT_mock_rep1_SPPE_TagDir | 43,702,697 |
| | | | WT_mock_rep2_SPPE_TagDir | 60,297,146 |
| | | WT 1 hour after 9J/m2 (n=2) | WT_UV1h_rep1_SSPE_TagDir | 62,042,662 |
| | | | WT_UV1h_rep2_SSPE_TagDir | 82,059,258 |
| | | WT 8 hours after 9J/m2 (n=2) | WT_UV_rep1_SPPE_TagDir | 52,616,690 |
| | | | WT_UV_rep2_SPPE_TagDir | 76,526,696 |
| | RPE1-iCas9 ELOF1-KO | ELOF1-KO input (n=2) | ELOF1_input_rep1_SPPE_TagDir | 40,511,010 |
| | | | ELOF1_input_rep2_SSPE_TagDir | 80,121,992 |
| | | ELOF1-KO mock-treated (n=2) | ELOF1_mock_rep1_SPPE_TagDir | 61,798,796 |
| | | | ELOF1_mock_rep2_SPPE_TagDir | 61,770,248 |
| | | ELOF1-KO 1 hour after 9J/m2 (n=2) | ELOF1_UV1hrs_rep1_SPPE_TagDir | 70,975,828 |
| | | | ELOF1_UV1hrs_rep2_SPPE_TagDir | 76,828,850 |
| | | ELOF1-KO 8 hours after 9J/m2 (n=2) | ELOF1_UV_rep1_SPPE_TagDir | 64,798,363 |
| | | | ELOF1_UV_rep2_SPPE_TagDir | 50,474,378 |
| | U2OS CSB-KO | CSB-KO mock-treated (n=2) | CSB-KO_mock_1 | 60,090,350 |
| | | | CSB-KO_mock_2 | 33,989,141 |
| | | CSB-KO 8 hours after 6J/m2 (n=2) | CSB-KO_UV_6J8h_1 | 55,433,652 |
| | | | CSB-KO_UV_6J8h_2 | 31,234,277 |
| | | CSB-KO 8 hours after 20J/m2 (n=1) | CSB-KO_UV_20J8h | 61,679,092 |
| RNA-seq | RPE1-iCas9 WT | WT mock-treated (n=3) | RNAseq_WT_Mock_rep1 | 16,570,265 |
| | | | RNAseq_WT_Mock_rep2 | 20,535,747 |
| | | | RNAseq_WT_Mock_rep3 | 20,356,970 |
| | | WT 24 hours after 9J/m2 (n=3) | RNAseq_WT_UV_rep1 | 30,231,046 |
| | | | RNAseq_WT_UV_rep2 | 18,430,293 |
| | | | RNAseq_WT_UV_rep3 | 22,828,990 |
| | RPE1-iCas9 ELOF1-KO | ELOF1-KO mock-treated (n=2) | RNAseq_ELOFKO_Mock_rep1 | 22,250,155 |
| | | | RNAseq_ELOFKO_Mock_rep2 | 18,412,158 |
| | | ELOF1-KO 24 hours after 9J/m2 (n=3) | RNAseq_ELOFKO_UV_rep1 | 33,575,741 |
| | | | RNAseq_ELOFKO_UV_rep2 | 20,292,635 |
| | | | RNAseq_ELOFKO_UV_rep3 | 22,376,948 |
| | RPE1-iCas9 CSB-KO | CSB-KO mock-treated (n=3) | RNAseq_CSBKO_Mock_rep1 | 26,689,782 |
| | | | RNAseq_CSBKO_Mock_rep2 | 23,198,147 |
| | | | RNAseq_CSBKO_Mock_rep3 | 20,027,395 |
| | | CSB-KO 24 hours after 9J/m2 (n=3) | RNAseq_CSBKO_UV_rep1 | 23,039,563 |
| | | | RNAseq_CSBKO_UV_rep2 | 12,906,509 |
| | | | RNAseq_CSBKO_UV_rep3 | 22,359,479 |

**Figure S1**

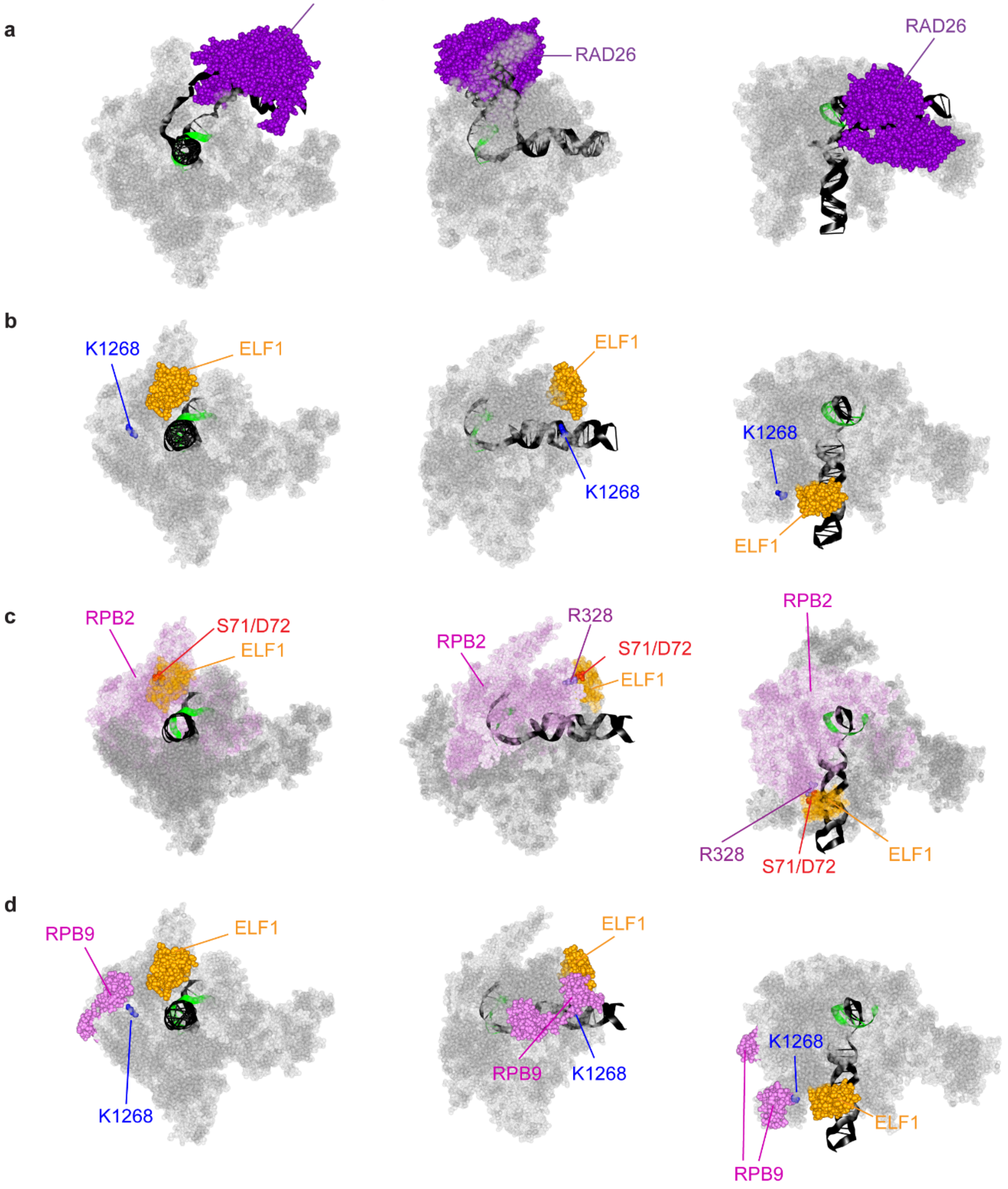
a
RAD26
RAD26
RAD26
b
K1268
ELF1
ELF1
K1268
K1268
ELF1
c
RPB2
S71/D72
ELF1
RPB2
R328
S71/D72
ELF1
RPB2
R328
S71/D72
ELF1
d
RPB9
ELF1
K1268
ELF1
RPB9
K1268
K1268
ELF1
RPB9

**Fig. S1. Structures of RNAPII bound to RAD26 or ELF1.** (**a-d**) Front-view, side-view and top-view of the following structures: (**a**) *S. cerevisiae* RAD26 (purple) bound to RNAPII (grey) (PBD: 5VVS). (**b**) *K. pastoralis* ELF1 (orange) bound to RNAPII (grey) with the RPB1-K1286 ubiquitylation site (K1264 in *K. pastoralis*) in blue. (PDB: 5XON). (**c**) *K. pastoralis* ELF1 (orange) bound to RNAPII (grey) (PDB: 5XON) with RPB2 in purple. Residues important for the ELF1(S71, D72) - RPB2 (R328) interaction are indicated. (**d**) *K. pastoralis* ELF1 (orange) bound to RNAPII (grey) with RPB9 in violet and the RPB1-K1286 ubiquitylation site (K1264 in *K. pastoralis*) in blue. (PDB: 5XON).

### Figure S2

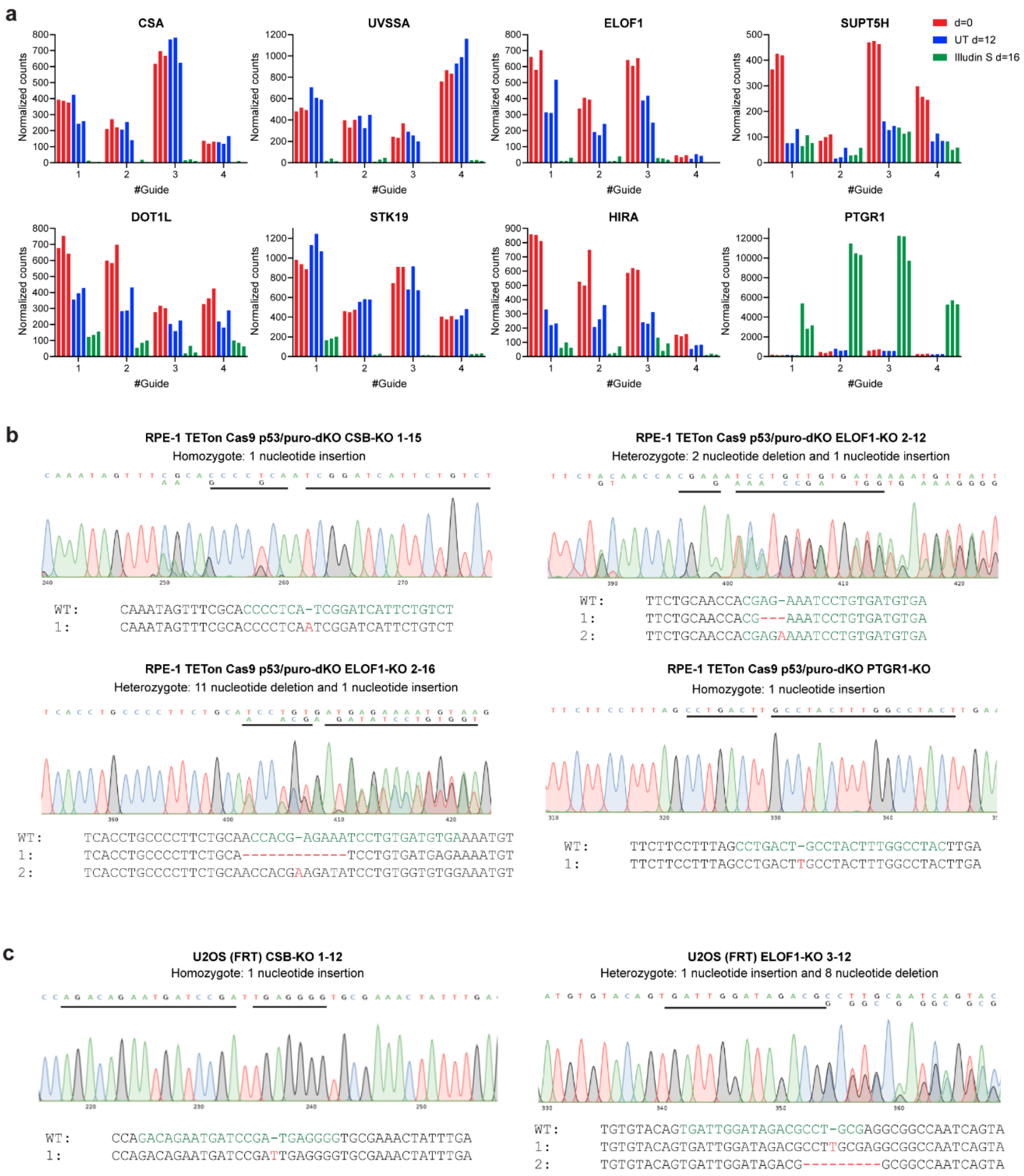
a
CSA
UVSSA
ELOF1
SUPT5H
DOT1L
STK19
HIRA
PTGR1
Normalized counts
#Guide
d=0
UT d=12
Illudin S d=16
b
RPE-1 TETon Cas9 p53/puro-dKO CSB-KO 1-15
Homozygote: 1 nucleotide insertion
WT: CAAATAGTTTCGCACCCCTCA-TCGGATCATTCTGTCT
1: CAAATAGTTTCGCACCCCTCAATCGGATCATTCTGTCT
RPE-1 TETon Cas9 p53/puro-dKO ELOF1-KO 2-12
Heterozygote: 2 nucleotide deletion and 1 nucleotide insertion
WT: TTCTGCAACCACGAG-AAATCCTGTGATGTGA
1: TTCTGCAACCACG---AAATCCTGTGATGTGA
2: TTCTGCAACCACGAGAAAATCCTGTGATGTGA
RPE-1 TETon Cas9 p53/puro-dKO ELOF1-KO 2-16
Heterozygote: 11 nucleotide deletion and 1 nucleotide insertion
WT: TCACCTGCCCCTTCTGCAACCACG-AGAAATCCTGTGATGTGAAAATGT
1: TCACCTGCCCCTTCTGCA------------TCCTGTGATGAGAAAATGT
2: TCACCTGCCCCTTCTGCAACCACGAAGATATCCTGTGGTGTGGAAATGT
RPE-1 TETon Cas9 p53/puro-dKO PTGR1-KO
Homozygote: 1 nucleotide insertion
WT: TTCTTCCTTTAGCCTGACT-GCCTACTTTGGCCTACTTGA
1: TTCTTCCTTTAGCCTGACTTGCCTACTTTGGCCTACTTGA
c
U2OS (FRT) CSB-KO 1-12
Homozygote: 1 nucleotide insertion
WT: CCAGACAGAATGATCCGA-TGAGGGGTGCGAAACTATTTGA
1: CCAGACAGAATGATCCGATTGAGGGGTGCGAAACTATTTGA
U2OS (FRT) ELOF1-KO 3-12
Heterozygote: 1 nucleotide insertion and 8 nucleotide deletion
WT: TGTGTACAGTGATTGGATAGACGCCT-GCGAGGCGGCCAATCAGTA
1: TGTGTACAGTGATTGGATAGACGCCTTGCGAGGCGGCCAATCAGTA
2: TGTGTACAGTGATTGGATAGACG---------GCGGCCAATCAGTA

**Fig. S2. Validation of knockout lines.** (**a**) Counts of the indicated sgRNAs from the CRISPR screen shown in Fig 1b. (**b**) Sanger sequencing of the indicated RPE1-iCas9 single *CSB*, *ELOF1*, or *PTGR1* knockout clones. (**c**) Sanger sequencing of the indicated U2OS (FRT) single *CSB*, or *ELOF1* knockout clones.

**Figure S3**

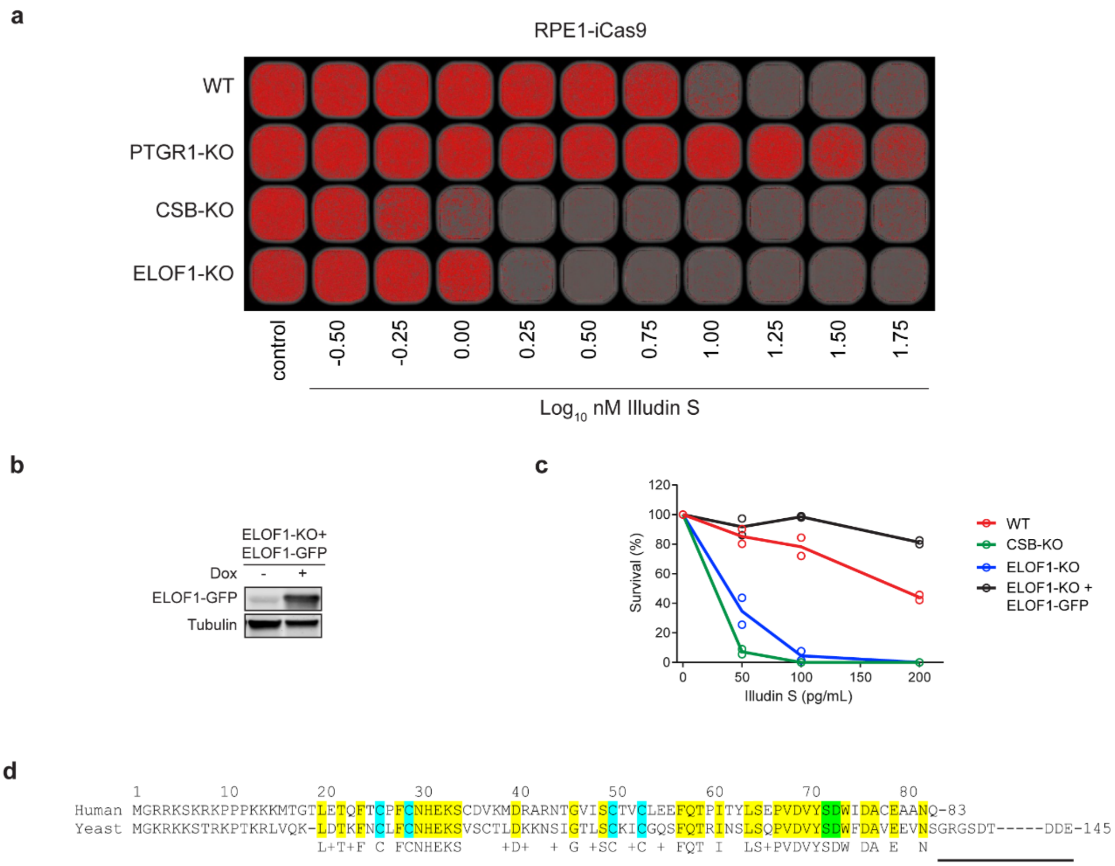
a
RPE1-iCas9
WT
PTGR1-KO
CSB-KO
ELOF1-KO
control
-0.50
-0.25
0.00
0.25
0.50
0.75
1.00
1.25
1.50
1.75
Log10 nM Illudin S
b
ELOF1-KO+
ELOF1-GFP
Dox
-
+
ELOF1-GFP
Tubulin
c
Survival (%)
120
100
80
60
40
20
0
0
50
100
150
200
Illudin S (pg/mL)
WT
CSB-KO
ELOF1-KO
ELOF1-KO +
ELOF1-GFP
d
1 10 20 30 40 50 60 70 80
Human MGRRKSKRKPPPKKKMTGTLETQFTCPFCNHEKSCDVKMDRARNTGVISCTVCLEEFQTPITYLSEPVDVYSDWIDACEAANQ-83
Yeast MGKRKKSTRKPTKRLVQK-LDTKFNCLFCNHEKSVSCTLDKKNSIGTLSCKICGQSFQTRINSLSQPVDVYSDWFDAVEEVNSGRGSDT-----DDE-145
L+T+F C FCNHEKS +D+ + G +SC +C + FQT I LS+PVDVYSDW DA E N

**Fig. S3. Human ELOF1 protects against Illudin S-induced toxicity** (**a**) Propidium-iodide (PI) Illudin S proliferation assay. (**b**) Western blot analysis of U2OS (FRT) *ELOF1*-KO cells complemented with inducible GFP-tagged ELOF1$^{WT}$. (**c**) Clonogenic Illudin S survival of U2OS (FRT) WT, *ELOF1*-KO, and *ELOF1*-KO complemented with ELOF1$^{WT}$-GFP. Each symbol represents the mean of an independent experiment (*n=2*), each containing 2 technical replicates. (**d**) Alignment of human ELOF1 and *S. cerevisiae* ELF1. Conserved residues are indicated in yellow, zinc-finger cysteines in magenta, residues involved in the RPB1 or RPB2 interaction in green. Note that the C-terminus of *S. cerevisiae* ELF1 (83-145) is absent in human ELOF1.

### Figure S4

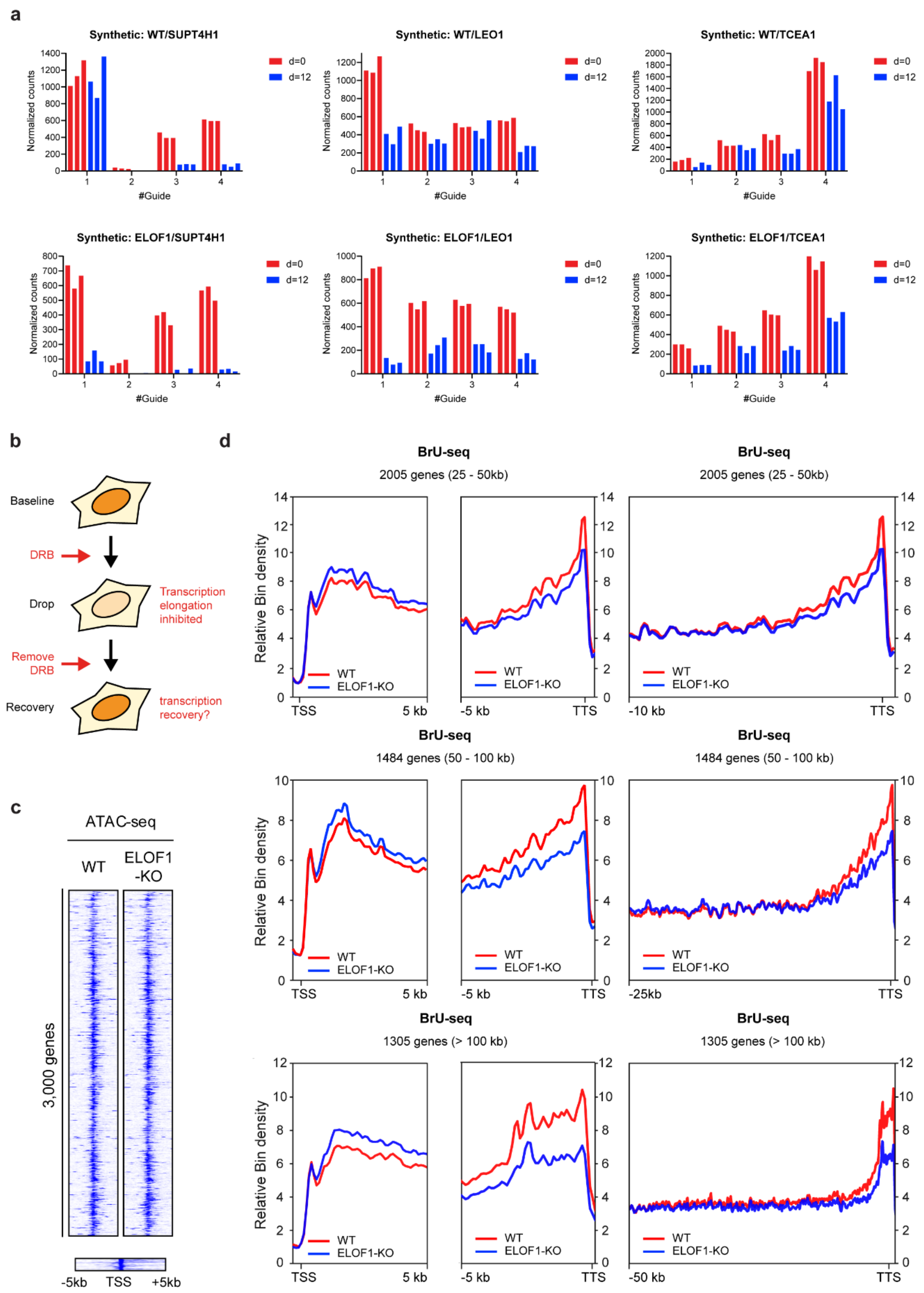
a
Synthetic: WT/SUPT4H1
Synthetic: WT/LEO1
Synthetic: WT/TCEA1
Synthetic: ELOF1/SUPT4H1
Synthetic: ELOF1/LEO1
Synthetic: ELOF1/TCEA1
Normalized counts
#Guide
d=0
d=12
b
Baseline
DRB
Drop
Transcription elongation inhibited
Remove DRB
Recovery
transcription recovery?
c
ATAC-seq
WT
ELOF1 -KO
3,000 genes
-5kb
TSS
+5kb
d
BrU-seq
2005 genes (25 - 50kb)
1484 genes (50 - 100 kb)
1305 genes (> 100 kb)
Relative Bin density
WT
ELOF1-KO
TSS
5 kb
-5 kb
TTS
-10 kb
-25kb
-50 kb

**Fig. S4. ELOF1 promotes transcription until the end of longer genes.** (**a**) Counts of the indicated sgRNAs from the isogenic cell-pair CRISPR screens shown in Fig 2a. (**b**) Experimental outline of the nascent transcription analysis after DRB wash-out. (**c**) Heatmaps of ATAC-seq data around the TSS (-5 kb until +5kb) of 3,000 genes of 3-100kb in unirradiated RPE1-iCas9 cells (WT or *ELOF1*-KO). (**d**) Metaplots of BrU signal (nascent transcription) in 2005 genes between 25-50 kb (upper), 1484 genes between 50-100 kb (middle), or 1305 genes of >100 kb in WT (red) or *ELOF1*-KO (blue) cells. BrU signal is shown in the first 5 kb after the TSS (left panel) and the last 5 kb before the TTS (middle panel).The right panel shows the BrU signal in the last 10 kb (25-50 kb genes), 25 kb (50-100 kb genes), or 50 kb (>100 kb genes) before the TTS.

### Figure S5

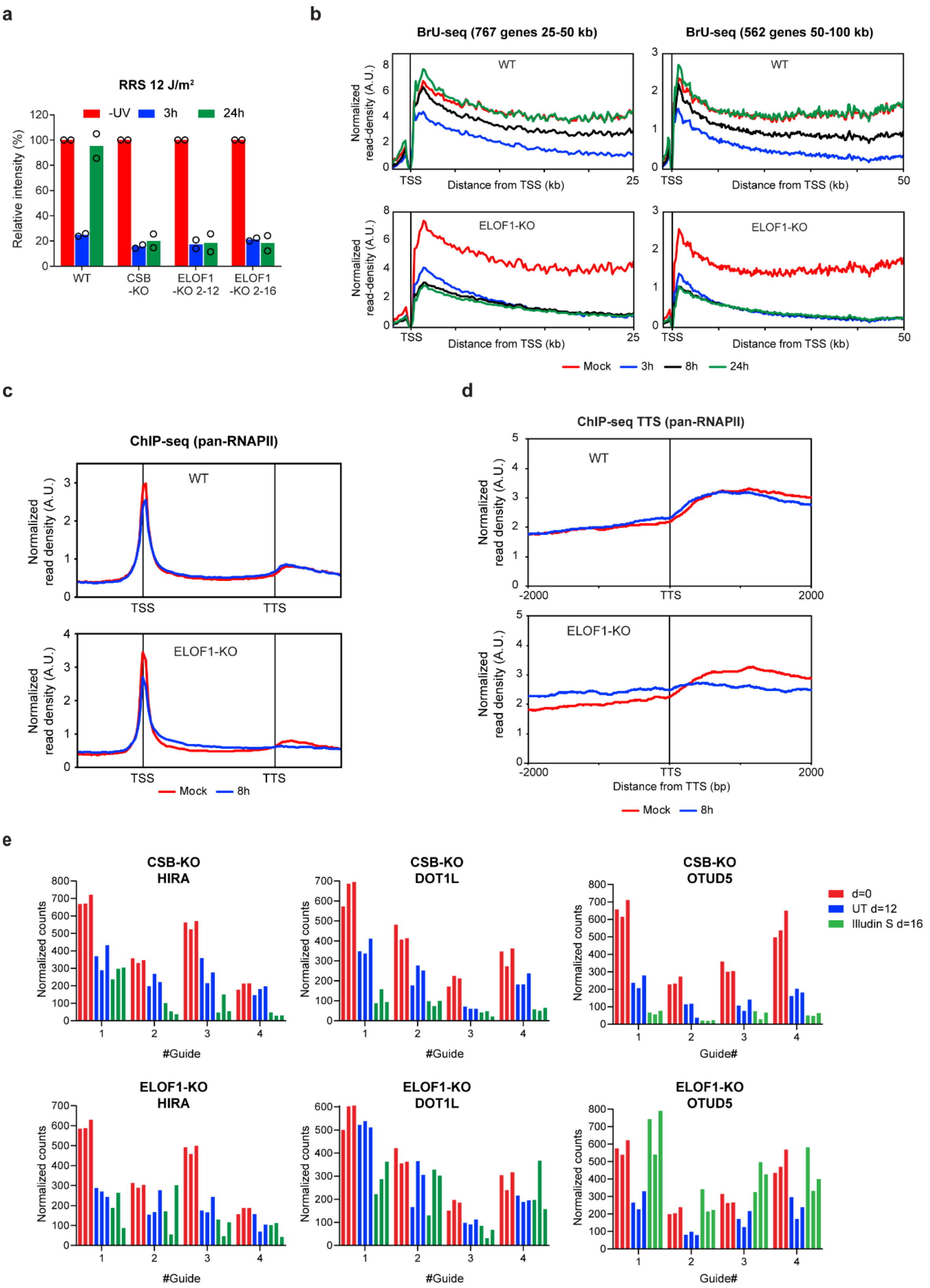
a
RRS 12 J/m²
-UV 3h 24h
Relative intensity (%)
WT
CSB -KO
ELOF1 -KO 2-12
ELOF1 -KO 2-16
b
BrU-seq (767 genes 25-50 kb)
BrU-seq (562 genes 50-100 kb)
WT
ELOF1-KO
Normalized read-density (A.U.)
Distance from TSS (kb)
Mock 3h 8h 24h
c
ChIP-seq (pan-RNAPII)
Normalized read density (A.U.)
WT
ELOF1-KO
TSS TTS
Mock 8h
d
ChIP-seq TTS (pan-RNAPII)
Normalized read density (A.U.)
WT
ELOF1-KO
Distance from TTS (bp)
Mock 8h
e
CSB-KO HIRA
CSB-KO DOT1L
CSB-KO OTUD5
ELOF1-KO HIRA
ELOF1-KO DOT1L
ELOF1-KO OTUD5
Normalized counts
#Guide
Guide#
d=0
UT d=12
Illudin S d=16

**Fig. S5. ELOF1 promotes genome-wide transcription recovery.** (**a**) Quantification of 5-EU levels in the indicated RPE1-iCas9 cells. Cells were either mock-treated (-UV) or UV-irradiated (3 h or 24 h; 12 J/m$^2$). Values were normalized to mock-treated levels for each cell-line. Each symbol represents the mean of an independent experiment (*n=2*) each containing 2 technical replicates, >80 cells collected per technical replicate. (**b**) Metaplots of BrU signal (nascent transcription) in 767 genes between 25-50 kb, or in 562 genes between 50-100 kb in WT (upper) or *ELOF1*-KO (lower) cells after mock treatment (red), or 3 h (blue), 8 h (black), and 24 h (green) after UV irradiation (9 J/m$^2$). (**c**) Averaged metaplots of pan-RNAPII ChIP-seq of 3,000 genes of 3-100kb from the TSS until the TTS in the indicated RPE1-iCas9 cells after mock-treatment (red) or at 8 h (blue) after UV irradiation (9 J/m2). (**d**) As in **c** showing a zoom around (-2 kb until +2 kb) the TTS. (**e**) Counts of the indicated sgRNAs from the isogenic cell-pair CRISPR screens shown in Fig 4a, b.

**Figure S6**

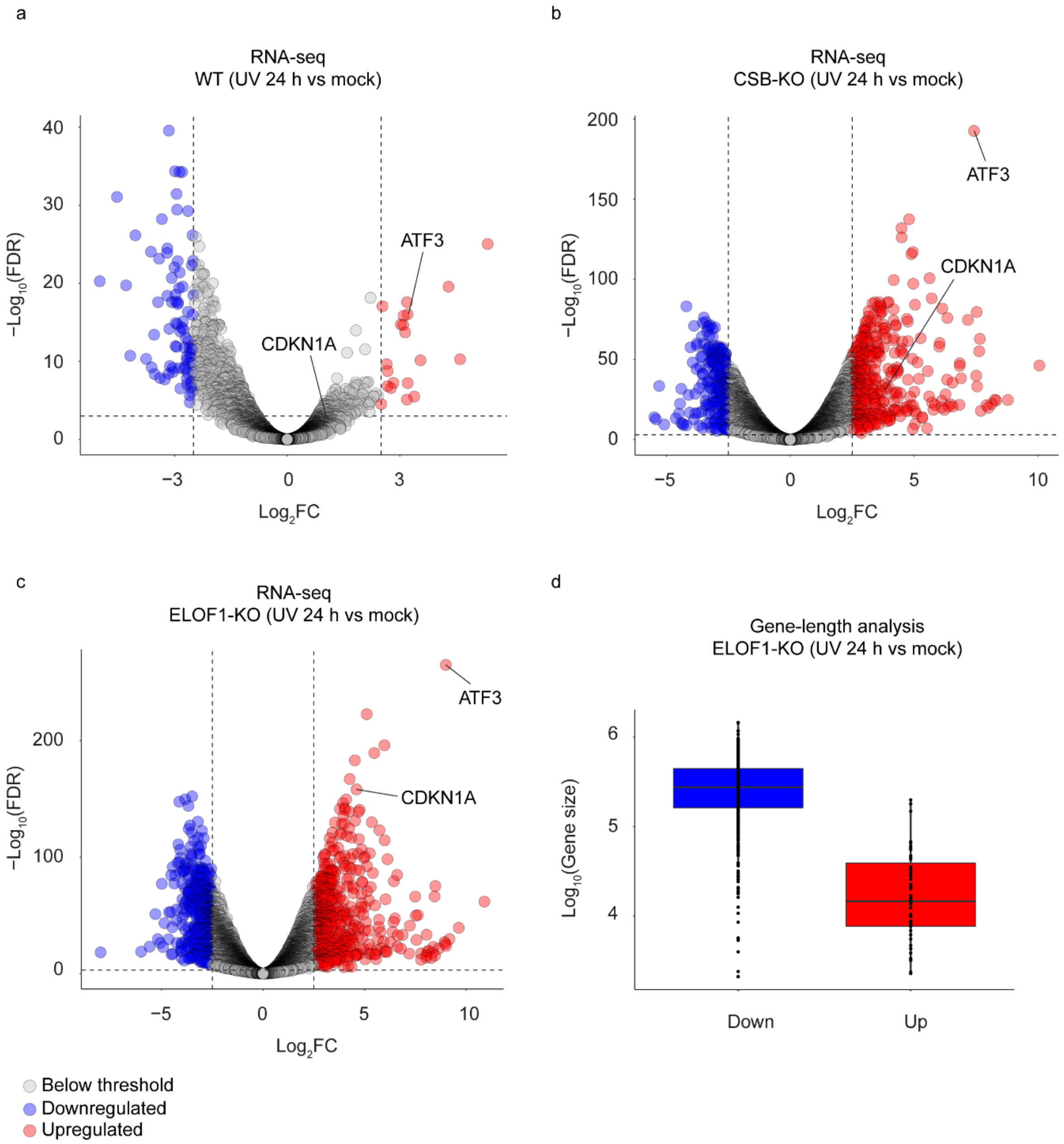
a
RNA-seq
WT (UV 24 h vs mock)
−Log10(FDR)
Log2FC
ATF3
CDKN1A
b
RNA-seq
CSB-KO (UV 24 h vs mock)
−Log10(FDR)
Log2FC
ATF3
CDKN1A
c
RNA-seq
ELOF1-KO (UV 24 h vs mock)
−Log10(FDR)
Log2FC
ATF3
CDKN1A
d
Gene-length analysis
ELOF1-KO (UV 24 h vs mock)
Log10(Gene size)
Down
Up
Below threshold
Downregulated
Upregulated

**Fig. S6. Global gene-expression changes in response to UV irradiation.** Volcano plots depicting the downregulation (in blue) or upregulation (in red) of gene expression in response to UV irradiation (9 J/m$^2$) after 24 h recovery. PolyA+ transcript levels were determined by RNA-seq and $\log_2$ fold-change was plotted against the significance ($-\log_{10}$ p-value) in the indicated RPE1-iCas9 cell lines: (**a**) WT, (**b**) *CSB*-KO, (**c**) *ELOF1*-KO. Only genes indicating at least 2 counts per million (CPM) in at least 33% of samples were included in the analysis. FDR-adjusted p-values < 0.05 were considered significant. Two short UV-response genes (*ATF3*, *CDKN1A*) are highlighted. Panel (**d**) shows the average length of the genes encoding either the 650 most significantly downregulated transcripts (in blue) or the 650 most significantly upregulated genes (in red).

**Figure S7**

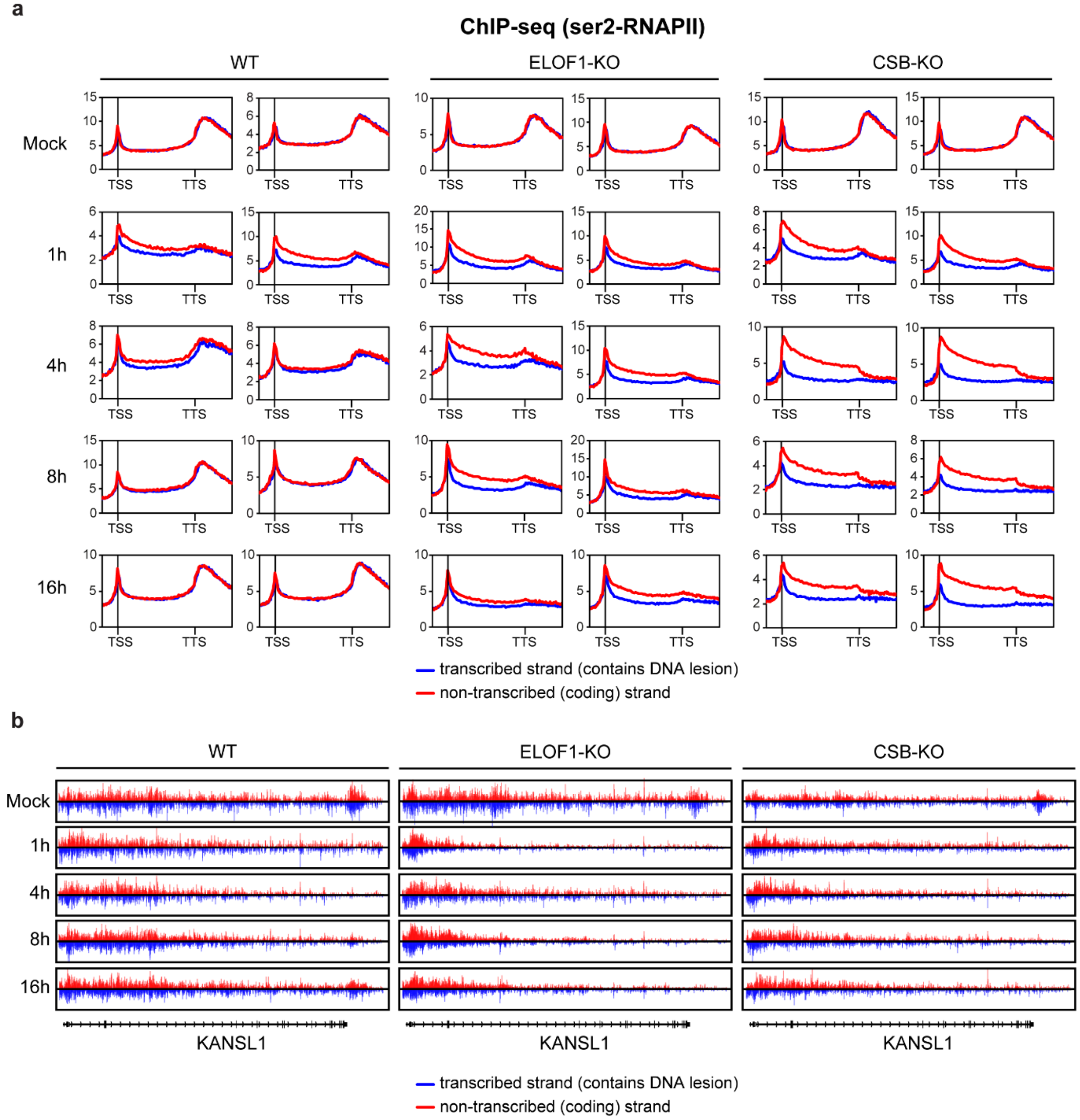
a
ChIP-seq (ser2-RNAPII)
WT
ELOF1-KO
CSB-KO
Mock
1h
4h
8h
16h
TSS
TTS
transcribed strand (contains DNA lesion)
non-transcribed (coding) strand
b
WT
ELOF1-KO
CSB-KO
Mock
1h
4h
8h
16h
KANSL1
transcribed strand (contains DNA lesion)
non-transcribed (coding) strand

**Fig. S7. TCR-seq using a Ser2-RNAPII antibody shows that ELOF1 is a core TCR factor.** (**a**) Individual metaplots (two replicates for each condition) of ser2-RNAPII TCR-seq of 3,000 genes for 3-100kb from the TSS until the TTS (-5kb and +5kb, respectively) in the indicated RPE1-iCas9 cells after mock-treatment or at 1 h, 4 h, or 8 h after UV irradiation (9 $J/m^2$). The coding (non-transcribed) strand in shown in red, while the template (transcribed) strand is shown in blue. (**b**) UCSC genome browser track showing the read density of TCR-seq data based on strand-specific ser2-RNAPII signal across the *KANSL1* gene after mock treatment, or at 1 h, 4 h, 8 h, or 16 h after UV irradiation (9 $J/m^2$) in WT, *CSB*-KO or *ELOF1*-KO cells. Reads from the coding (non-transcribed) strand are shown in red, while reads from the template (transcribed) strand are shown in blue.

**Figure S8**

**a**

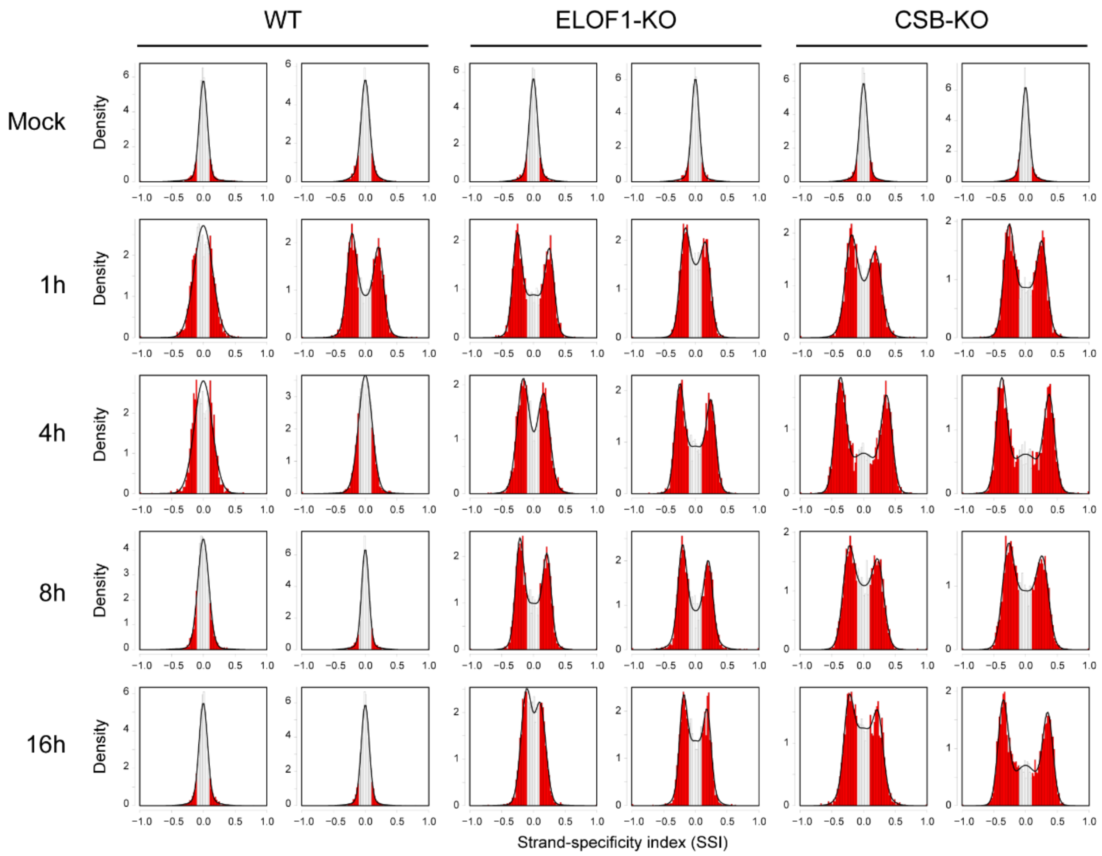
WT
ELOF1-KO
CSB-KO
Mock
1h
4h
8h
16h
Density
Strand-specificity index (SSI)

**b**

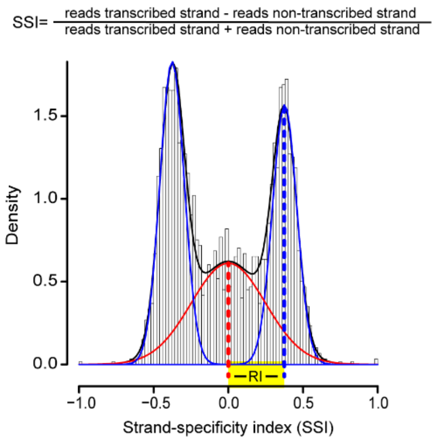
SSI= reads transcribed strand - reads non-transcribed strand / reads transcribed strand + reads non-transcribed strand
Density
RI
Strand-specificity index (SSI)

**Fig. S8. Histogram plots of the strand-specificity index.** (**a**) Frequency distribution plots of the gene-by-gene ser2-RNAPII strand-specificity index (SSI). SSIs below -0.1 or above 0.1 are presented in red. A unimodal distribution indicates no strand-bias (and thus no DNA damage in the template strand), while a trimodal distribution reflects a strand-bias caused by DNA damage in the template strand. (**b**) Explanation of the recovery index (RI) calculation. Per sample, a frequency distribution plot was generated of the per-gene strand specificity index (SSI; defined as the relative difference in read density between the transcribed and non-transcribed strands) of 3,000 genes of 3-100kb. The RI is subsequently obtained by fitting a mixture of 3 Gaussian distributions, corresponding to the undamaged gene fraction (SSI=0) and two unrepaired gene fractions (i.e. |SSI|<>0).

**Figure S9**

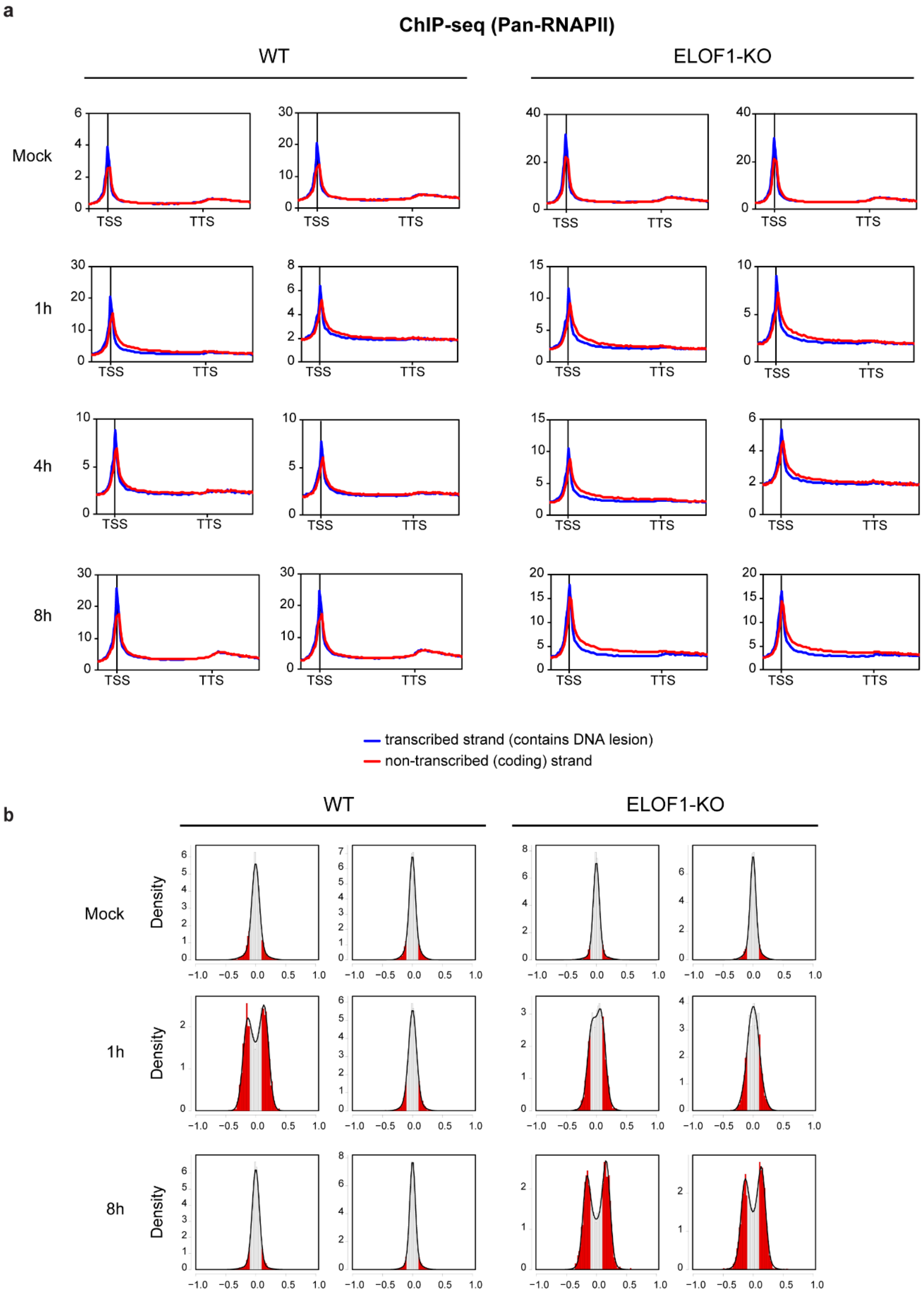
a
ChIP-seq (Pan-RNAPII)
WT
ELOF1-KO
Mock
1h
4h
8h
TSS
TTS
transcribed strand (contains DNA lesion)
non-transcribed (coding) strand
b
WT
ELOF1-KO
Mock
1h
8h
Density

**Fig. S9. Validation of TCR-seq with a pan-RNAPII antibody.** (**a**) Individual metaplots (two replicates for each condition) of pan-RNAPII TCR-seq of 3,000 genes of 3-100kb from the TSS until the TTS (-5kb and +5kb, respectively) in the indicated RPE1-iCas9 cells after mock-treatment or at 1 h, 4 h, or 8 h after UV irradiation (9 J/$m^2$). The coding (non-transcribed) strand in shown in red, while the template (transcribed) strand is shown in blue. (**b**) Frequency distribution plots of the gene-by-gene ser2-RNAPII strand-specificity index (SSI). SSIs below -0.1 or above 0.1 are presented in red. A unimodal distribution indicates no strand-bias (and thus no DNA damage in the template strand), while a trimodal distribution reflects a strand-bias caused by DNA damage in the template strand.

**Figure S10**

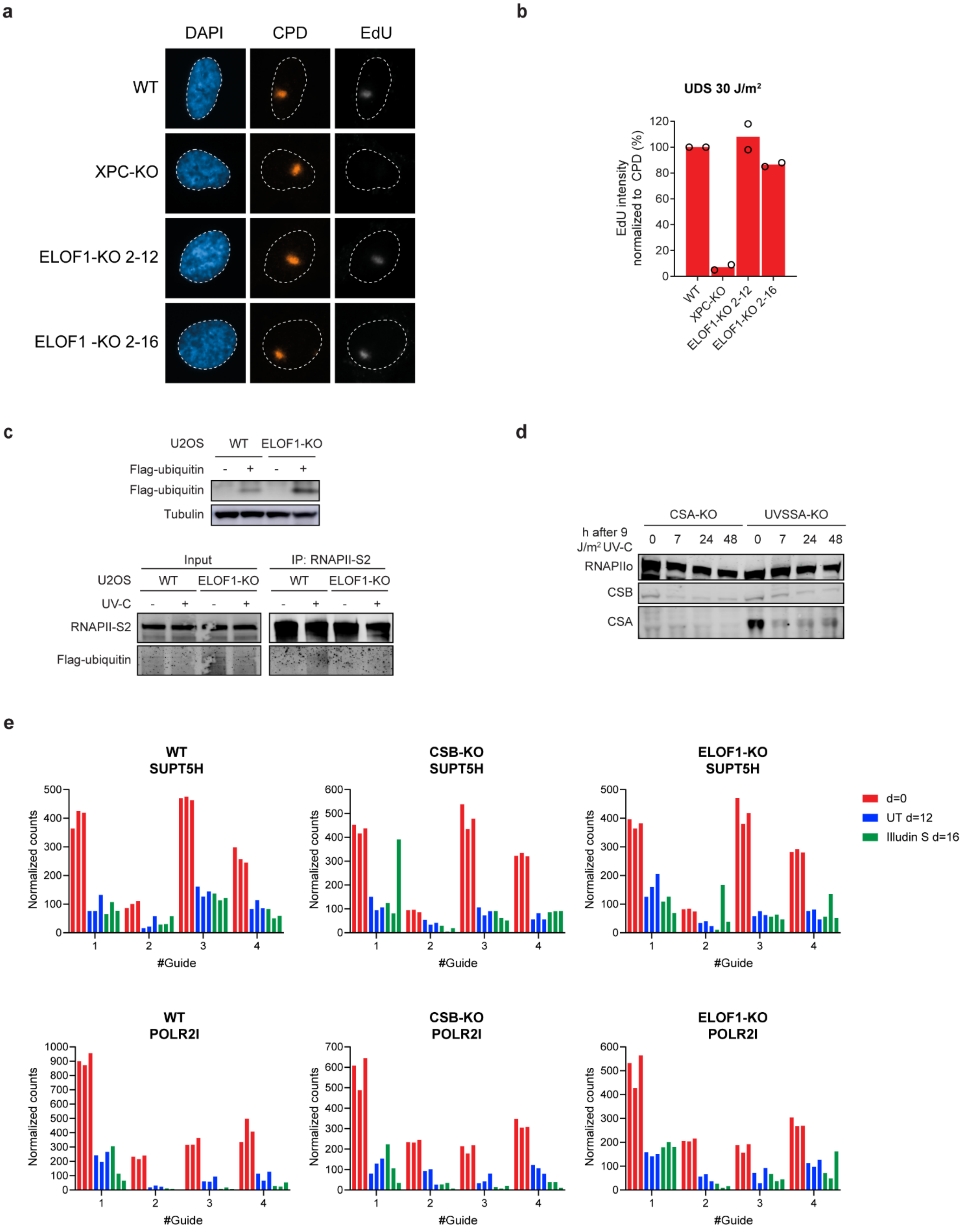
a
DAPI
CPD
EdU
WT
XPC-KO
ELOF1-KO 2-12
ELOF1 -KO 2-16
b
UDS 30 J/m²
EdU intensity normalized to CPD (%)
0
20
40
60
80
100
120
WT
XPC-KO
ELOF1-KO 2-12
ELOF1-KO 2-16
c
U2OS
WT
ELOF1-KO
Flag-ubiquitin
- + - +
Flag-ubiquitin
Tubulin
Input
IP: RNAPII-S2
U2OS
WT
ELOF1-KO
WT
ELOF1-KO
UV-C
- + - + - + - +
RNAPII-S2
Flag-ubiquitin
d
CSA-KO
UVSSA-KO
h after 9 J/m² UV-C
0 7 24 48 0 7 24 48
RNAPIIo
CSB
CSA
e
WT SUPT5H
CSB-KO SUPT5H
ELOF1-KO SUPT5H
Normalized counts
#Guide
1
2
3
4
d=0
UT d=12
Illudin S d=16
WT POLR2I
CSB-KO POLR2I
ELOF1-KO POLR2I
Normalized counts
#Guide

**Fig. S10. ELOF1 is not involved in global genome repair.** (**a**) Unscheduled DNA synthesis (UDS) in the indicated RPE1-iCas9 lines (WT, *XPC*-KO, *ELOF1*-KO) after local irradiation through a 5 µm millipore filter with 30 J/m$^2$, and subsequently labeled with EdU to monitor global genome repair (GGR) activity. DNA damage was identified by CPD staining. (**b**) Quantification of EdU levels normalized to CPD signal. Each symbol represents the mean of an independent experiment (*n=2*), each containing 2 technical replicates, >80 cells collected per technical replicate (**c**) Western blot analysis (upper panel) and IP on RNAPII (lower panel) of U2OS (FRT) WT and *ELOF1*-KO cells stably expressing Flag-ubiquitin. (**d**) Western blot analysis of CSA protein levels in the indicated RPE1-iCas9 cells after mock treatment, or 7h, 24h, and 48h after UV irradiation (9 J/m$^2$). (**e**) Counts of the indicated sgRNAs from the isogenic cell-pair CRISPR screens shown in Fig 4a, b.
